# Mechanics-dependent Global Nuclear Eviction and Site-Specific Recruitment of YAP Regulates DNA Damage Responses

**DOI:** 10.1101/2025.11.23.690063

**Authors:** Shantam Yagnik, Aprotim Mazumder

## Abstract

Yes-associated protein (YAP), a transcriptional coactivator, plays key roles in cell growth, proliferation and apoptosis, and its levels are frequently dysregulated in cancers. YAP levels in the nucleus are highly sensitive to nuclear mechanical cues, and such cues are also parallelly emerging to be a key modulator of DNA Damage Responses (DDR). However, whether DNA-damage can induce mechanical changes that regulate downstream events such as YAP localization and that in turn feeds back onto DDR activation, remains unknown. In this study, we report that YAP translocates in a nuclear mechanics-dependent manner upon induction of Double Strand Breaks (DSBs). This translocation is not a mere epiphenomenon, and we find that: first, global nuclear eviction of YAP enhances DDR signaling; second, local enrichment of YAP at DNA damage sites promotes recruitment of DNA repair proteins previously identified as potential interactors of YAP or its partner TEAD1. Together, these findings indicate that YAP is not only a transcriptional coactivator, but also plays an under-appreciated role in regulating DDR.

## Introduction

Yes-associated protein (YAP), a transcriptional coactivator, plays a crucial role in controlling cell proliferation, development and organ size [1, 2]. Elevated levels and hyperactivation of YAP have been observed in several cancers [3]. It exists in humans in two isoforms, YAP1 and YAP2, which differ from each other by the presence of a second WW domain in YAP2. YAP functions by binding to transcription factors such as TEAD [4–6], RUNX1/2 [7], and p73 [8], thus modulating gene transcription and bringing about different functional outcomes. YAP is a downstream target of the Hippo pathway, whose primary function is to regulate the activity of YAP [9]. The Hippo pathway does so through multiple Ste20 kinase family proteins (MST1/2, TAO family kinases (TAO) and several MAP4K members) [10, 11]. These kinases, along with accessory proteins like SAV1, MOB1, NF2 phosphorylate and activate the LATS1 and LATS2 (LATS 1/2) kinases. Activated LATS1/2 phosphorylate YAP at S127, creating a 14-3-3 binding site that promotes cytoplasmic retention of YAP and decreases its nuclear activity [12, 13]. Recent studies have, however, shown that YAP can be regulated in the nucleus independent of the Hippo pathway. In particular, mechanical cues can influence the nuclear localization of YAP. For example, Mesenchymal Stem Cells (MSCs), depend on YAP regulation to differentiate into osteogenic or adipogenic lineages. This regulation of YAP was shown to be controlled by the substrate stiffness the cells were exposed to [14, 15], positioning YAP as a key mediator of mechanical signals to the nucleus. Focal adhesions, cell-cell junctions communicate with the nuclear membrane to form a bridge to both sense and respond to internally or externally generated forces [16–18]. The actin cytoskeleton is one of the main components that can sense and respond to these forces experienced by the cell [19]. YAP nuclear localization was observed to be sensitive to changes in actin levels, particularly F-Actin, through both Hippo (LATS1/2) dependent and independent mechanisms [20, 21]. Tension sensing by focal adhesions have been shown to increase cell spreading and increased YAP/TAZ activity [22, 23]. Cell spreading and subsequent nuclear flattening, dependent on focal adhesions, has been found to promote nuclear levels of YAP. This is attributed to size and mechano-stability of YAP, which permit its differential translocation across nuclear pores whose permeability increases during nuclear flattening [24]. Additionally, F-Actin regulates YAP activity through competition with ARID1A, a component of the SWI/SNF chromatin remodelling complex. YAP binds to ARID1A, which blocks its interaction with TEAD. When nuclear F-Actin levels increase, such as in response to mechanical cues like an increase in adhesive area, it causes ARID1A to associate more with F-Actin instead of YAP, thus freeing YAP to associate with TEAD and activate transcription [25].

Parallel to the studies on the effects of mechanics on YAP localization, a growing number of studies are also revealing a connection between nuclear mechanics and DNA Damage Responses (DDR) [26]. DDR is largely mediated by signaling cascades driven by phosphorylation, initiated by the upstream kinases like ATM, ATR, and DNA-PK, which orchestrate a broad network of cellular processes to maintain genomic integrity. Recent studies have observed changes in nuclear mechanics on DNA damage [26], with both ATM and ATR shown to be sensitive to nuclear mechanical forces. ATM was shown to control nuclear stiffness by modulating Lamin A levels [27] and ATR was shown to directly sense mechanical forces along the nuclear envelope [28]. While the effects of mechanical perturbations on DDR kinases are increasingly appreciated, the converse connection between DNA damage and nuclear mechanics remain more tenuous.

In response to genotoxic stresses caused by cisplatin, it was shown that phosphorylation of YAP at S127 by Akt causes its cytoplasmic retention and spatial separation from nuclear transcription factors [29]. Further, YAP has been identified to be a downstream target of ATM/ATR kinases, mainly through two mechanisms. First, double-strand DNA breaks activate ATM, which phosphorylates RASSF1A at Serine 131, leading to its dimerization and binding to MST2 kinase. This enhances MST kinase activity, phosphorylating LATS1/2, which then phosphorylates YAP at S127, promoting YAP-p73 association and shifting YAP function from oncogenic to tumour-suppressive [30–32]. Second, ATM also phosphorylates cAbl, which in turn binds and phosphorylate YAP at Tyr 357 residue, stabilizing the YAP-p73 complex [33–36].

In light of the studies discussed above, and given the emerging role of nuclear mechanics in regulating YAP nuclear localization, we sought to investigate whether DNA damage can induce changes in nuclear mechanics, influence YAP levels in the nucleus and downstream function. We also investigate whether such potential relocalization is an epiphenomenon of changes in cellular mechanics, or can it have a potential bearing on the damage response through ATM/ATR signaling. We find that YAP is a mediator between changes in nuclear mechanics and DNA damage responses, with its levels having a direct effect on DDR. We also find that while YAP is globally depleted from nuclei in response to Double Strand Break (DSB) induction, it is intriguingly recruited to the site of DNA damage and has a potential role in DDR. DDR and YAP-dependent programs are independently known to be gated by cell mechanics; in this study we show that the three are functionally interconnected.

## Results

### YAP is evicted from the nucleus on DNA damage independent of Hippo pathway and is enriched at the site of DNA damage

ATM/ATR activity is well documented in the case of DSB responses [37–39]. To investigate if DDR has a role in regulating the levels of YAP upon DNA damage, we adopted a system of Hoechst sensitization mediated DSB induction using a 405nm laser [40, 41], followed by live tracking of cells over time. (Fig. 1A). Hoechst-sensitized U2OS cells expressing EGFP-YAP1 (YAP1 fused with the Enhanced Green Fluorescent Protein) were irradiated using a point ROI with a 405nm laser and followed over time. EGFP-YAP1 was found to exit the nucleus over minutes after irradiation. To quantify this eviction, we quantified the EGFP-YAP1 nuclear to cytoplasmic intensity ratio (hereafter referred to as YAP ratio) over time (Fig. 1B, C) (see Materials and Methods for details). A decreasing ratio is indicative of cytoplasmic translocation of YAP. We performed a second control experiment comparing Hoechst-stained and non-Hoechst stained EGFP-YAP1 expressing cells subjected to identical 405 nm laser micro-irradiation. YAP localization was followed over time under both conditions. Importantly, no decrease in the nuclear-to-cytoplasmic YAP ratio was observed in non-Hoechst stained cells despite identical irradiation conditions, whereas Hoechst-stained cells showed a robust reduction in nuclear YAP following irradiation (Supp. Fig. 1 A, B). Thus, YAP eviction follows DNA damage upon irradiation of Hoechst-sensitized cells. We also addressed if the observed nuclear loss is specific to YAP. To answer this, we examined the behaviour of an unrelated nuclear protein under identical experimental conditions. Specifically, we used EGFP-Nucleolin, a primarily nucleolar protein tagged with the same EGFP similar to the EGFP-YAP1 construct, which shows both nuclear and nucleolar expression. Nucleolin is known to translocate from the nucleolus to the nucleoplasm upon damage [42] but is not expected to be evicted from the nucleus. In contrast to EGFP-YAP1, we did not observe any significant decrease in the nuclear-to-cytoplasmic ratio of EGFP-Nucleolin following DNA damage compared to undamaged cells (Supp. Fig. 1 C, D). These experiments used EGFP-YAP1 transfected cells, enabling live-cell dynamics, but could have potential overexpression artefacts. So, we attempted to capture the same phenomenon of YAP translocation using antibody staining for the endogenous protein. Since, YAP ratio showed maximum decline at 20mins post irradiation with EGFP-YAP1, we chose this time point after damage to perform immunofluorescence (IF) (Fig. 1D). Indeed, we saw a similar decline (∼30%) of YAP ratio at this time point as we see in live cells with EGFP-YAP1 on DNA damage (Fig. 1 E, F). Moreover, this reduction was dose-dependent, with greater YAP eviction observed at higher laser intensities (2% vs. 20%) (Supp. Fig. 2A, B). The increase in γH2AX intensity at increasing doses further confirmed the graded DNA damage response (Supp. Fig. 2A). Thus, even endogenous YAP1 translocates upon laser-induced DSB induction. Additionally, beyond the global nuclear eviction of YAP, rather intriguingly, we also observed local enrichment of YAP at the sites of DNA damage/laser-irradiation in these IF experiments (Fig. 1 E), which was also observed at lower (2%) damage (Supp. Fig 2 A). Why a transcriptional coactivator should accumulate at sites of damage was not immediately obvious. To test whether this phenomenon is conserved in an adherent cell-line from a different species, we examined endogenous YAP localization in the mouse fibroblast cell line NIH/3T3 upon identical DNA damage conditions. Although NIH/3T3 cells display a lower basal nuclear YAP ratio compared to U2OS cells under control conditions, DNA damage still produced a small but clear and reproducible decrease in nuclear YAP levels relative to undamaged controls (Supp. Fig. 2 D, E). Thus, the damage-induced reduction in nuclear YAP is conserved despite differences in baseline YAP localization between the two cell lines. We also tested YAP levels upon damage with Neocarzinostatin (NCS), which induces DSBs without the need for photosensitizing dyes. We observe a modest but consistent and statistically significant decrease in nuclear YAP levels across independent experimental replicates (Supp. Fig. 2 F, G), indicating that YAP eviction may be a general feature of DSB response rather than being specific to laser-induced clustered DSBs. While globally the nuclear YAP concentrations decreased in the laser irradiation experiments (as indicated by a lower nuclear to cytoplasmic ratio), locally there was an enrichment of YAP at the site of damage (Fig. 1 G; Supp. Fig 2 C). This may hint at a different role of YAP in affecting DDR apart from its global nuclear eviction.

**Figure 1.**
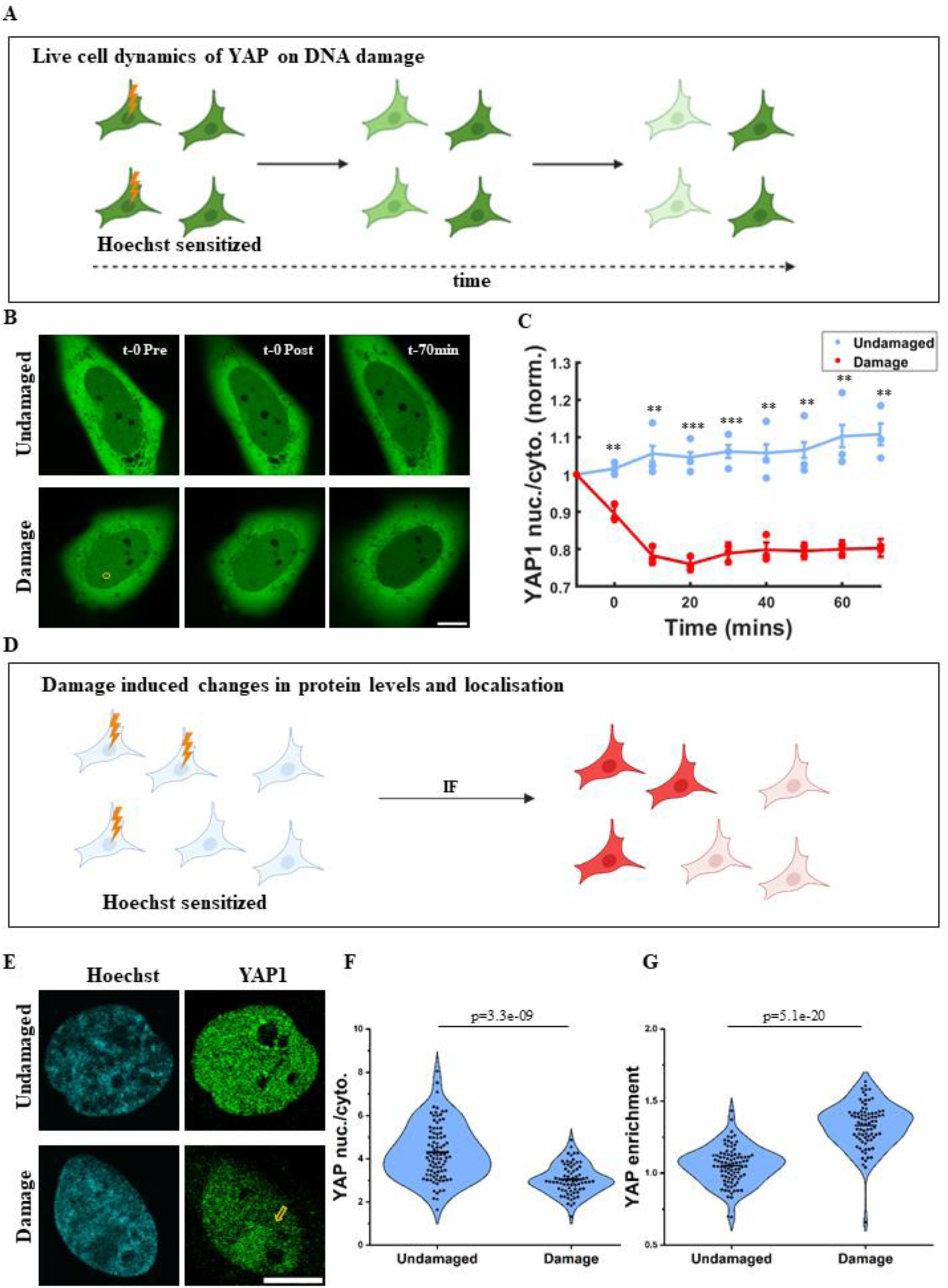
YAP is evicted from the nucleus on DNA damage and is localised at the site of DNA damage. (A) Schematic illustrating irradiation of Hoechst sensitized cells overexpressing EGFP-YAP1 with a 405nm laser to follow the dynamics of YAP over time. (B) Representative images showing dynamics of EGFP-YAP1 at different time points in U2OS cells. The top panel shows undamaged cells and the bottom panel shows cells just before causing damage (t-0 Pre), just after damage (t-0 Post) and 70min after damage. The yellow circle represents the ROI of laser irradiation in the bottom panel. The images are displayed as average z-projections. The scale bar is 10 microns. (C) Timelapse curves showing the dynamics of EGFP-YAP1 at different time points, in undamaged control (blue) and damaged cells (red). The nuclear-to-cytoplasmic intensity ratio of YAP1 was measured at each time point and normalised to the initial value. Each dot represents the experiment-wise mean YAP ratio. The curve shows the overall mean ± SEM from three independent experiments (N=3, n>15 cells per condition). Statistics: The stars denote the p-values for each time point calculated using the Student’s t-test (p>0.05-ns (non-significant), 0.01<p<0.05 - *, 0.001<p<0.01 - **, p<0.001 - ***). (D) Schematic illustrating irradiation of Hoechst sensitized cells with a 405nm laser followed by immunofluorescence-based labelling of protein of interest to capture changes in intensity of endogenous protein and compare them with undamaged control in the same plate. (E) Representative images showing U2OS cell nuclei with Hoechst (cyan), intensity of endogenous YAP1 levels (green) in undamaged cells (top panel) and laser irradiated damaged cells (bottom panel). The yellow circle represents the ROI of laser irradiation in the bottom panel. The yellow arrow indicates the enrichment of YAP1 after damage. The scale bar is 10 microns. (F) Violin plot with distribution quantifying nuclear to cytoplasmic intensity ratio of YAP1 (YAP1 nuc./cyto. or YAP ratio) along with the mean in undamaged and damaged cells. Each dot is the YAP ratio from a single cell (N=3, n>70 cells per condition). The p-values from the distributions are calculated using the Kolmogorov-Smirnov test. (G) Violin plot with distribution quantifying YAP1 enrichment along with the mean in undamaged and damaged cells. Each dot is the YAP enrichment from a single cell (N=3, n>70 cells per condition). Enrichment is defined as the ratio of mean intensity at the site of damage to the mean intensity in the rest of the nucleus. For undamaged cells a random spot in the nucleus for each cell was chosen from the Hoechst channel as ROI. The p-values from the distributions are calculated using the Kolmogorov-Smirnov test.

**Figure 2.**
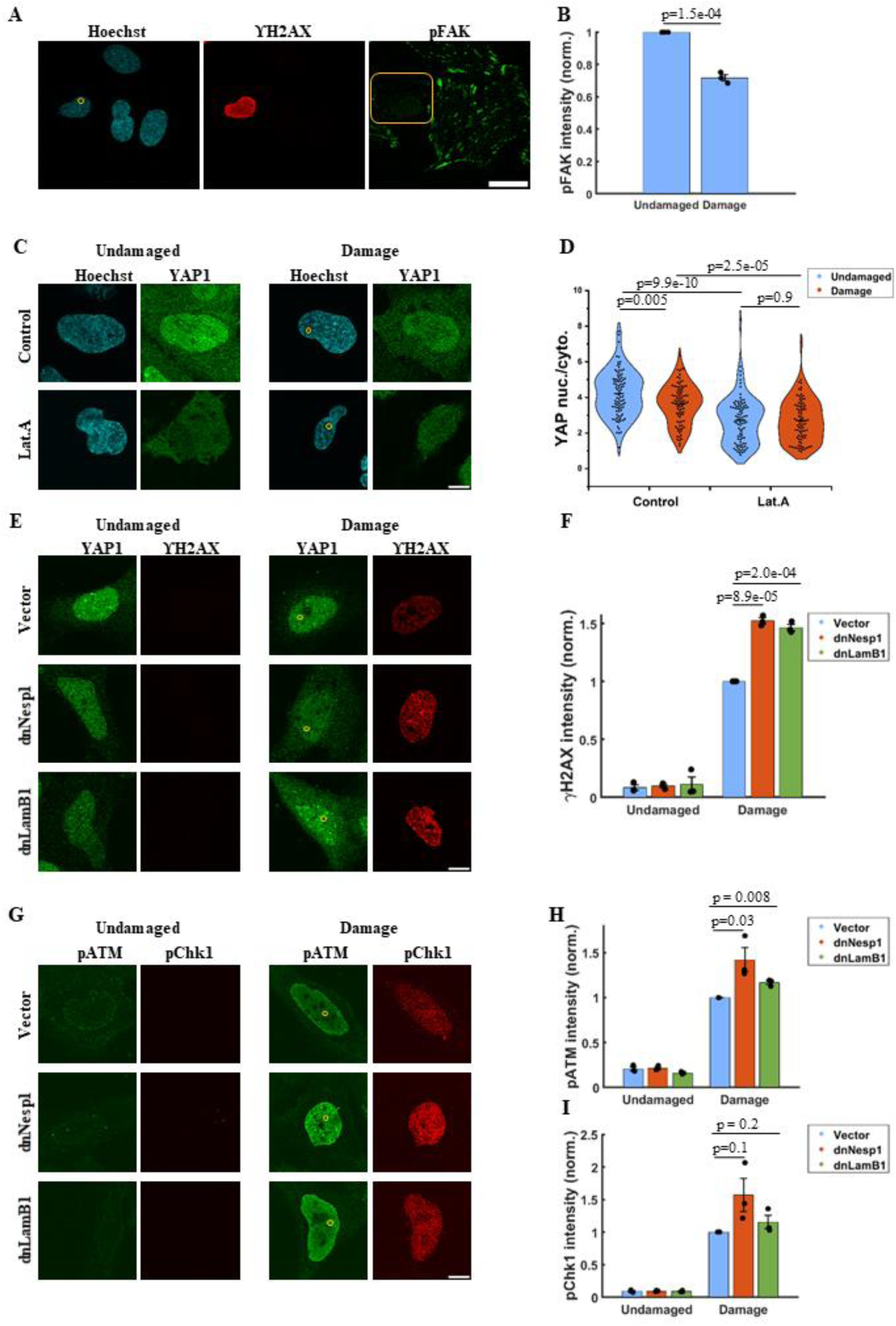
Nuclear mechanical changes on DNA damage affect YAP translocation and damage responses. (A) Representative images showing U2OS cell nuclei with Hoechst (cyan), intensity of endogenous ϒH2AX (red) and pFAK (pY397) (green). The yellow circle represents the ROI of laser irradiation in the Hoechst channel. The images are maximum intensity projected. The scale bar is 40 microns. (B) Bar graph quantifying normalised total pFAK intensity at focal adhesions in undamaged versus damaged U2OS cells. Each dot represents the experiment-wise mean, normalised to the corresponding undamaged control, and pooled across experiments (N=3, n>70 cells per condition). The p-values from the distributions are calculated using unpaired Student’s t-test. (C) Representative images showing U2OS cell nuclei with Hoechst (cyan), intensity of endogenous YAP1 (green) in undamaged cells (left panel) and damaged cells (right panel) in both vehicle treated (Control, DMSO, top) or Latrunculin A treated (Lat.A, bottom) cells. The yellow circle represents the ROI of laser irradiation in the right panel. The scale bar is 10 microns. (D) Violin plot with distribution quantifying nuclear to cytoplasmic intensity ratio of YAP1 (YAP1 nuc./cyto. or YAP ratio) in undamaged and damaged cells of both vehicle treated (Control, DMSO, blue) or Latrunculin A treated (Lat.A, orange) cells. Each dot is the YAP ratio from a single cell (N=3, n>70 cells per condition). The p-values from the distributions are calculated using the Kolmogorov-Smirnov test. (E) Representative images showing intensities of endogenous YAP1 (green) and ϒH2AX (red) in undamaged cells (left panel) and damaged cells (right panel) overexpressing mApple vector control (Vector, top), dominant negative Nesprin1 (dnNesp1, middle) and dominant negative Lamin B1 (dnLamB1, bottom) respectively. The yellow circle represents the ROI of laser irradiation in the right panel. The scale bar is 10 microns. (F) Bar graph quantifying intensity of ϒH2AX in C1-mApple (Vector, blue), dnNesp1 (orange) and dnLamB1 (green) overexpressing U20S cells undamaged and damaged conditions. Each dot represents the experiment-wise mean, normalised to the corresponding damaged control (mApple), and pooled across experiments (N=3, n>70 cells per condition). Normalisation is done to the damaged mApple control, as all the undamaged samples have very low levels of staining making normalization to those difficult. The p-values from the distributions are calculated using unpaired Student’s t-test. (G) Representative images showing intensities of endogenous pATM (pS1981) (green) and pChk1 (pS345) (red) in undamaged cells (left panel) and damaged cells (right panel) overexpressing mApple vector control (Vector, top), dominant negative Nesprin1 (dnNesp1, middle) and dominant negative Lamin B1 (dnLamB1, bottom). The yellow circle represents the ROI of laser irradiation in the right panel. The scale bar is 10 microns. (H) Bar graph quantifying intensity of pATM in C1-mApple (Vector, blue), dnNesp1 (orange) and dnLamB1 (green) overexpressing U20S cells undamaged and damaged conditions. Each dot represents the experiment-wise mean, normalised to the corresponding damaged control, and pooled across experiments (N=3, n>70 cells per condition). The p-values from the distributions are calculated using unpaired Student’s t-test. (I) Bar graph quantifying intensity of pChk1in C1-mApple (Vector, blue), dnNesp1 (orange) and dnLamB1 (green) overexpressing U20S cells undamaged and damaged conditions. Each dot represents the experiment-wise mean, normalised to the corresponding damaged control, and pooled across experiments (N=3, n>70 cells per condition). The p-values from the distributions are calculated using unpaired Student’s t-test.

Since YAP/TAZ are the main transducers of the Hippo pathway, we next investigated whether the decrease in YAP nuclear levels at this time point is dependent on the Hippo pathway. YAP phosphorylation on specific serine residues, especially at the S127 by LATS1/2 targets it for proteasomal degradation. For both YAP1 and YAP2 this phosphorylation on S127 has been shown to be a factor promoting cytoplasmic translocation and S127A mutant that cannot be phosphorylated has been used to underscore this point [12, 13, 43]. To test whether S127 phosphorylation is important for cytoplasmic translocation, we performed live cell experiments in cells expressing WT YAP2 and S127A YAP2 with DsRed tags as previously described [13]. YAP2 too evidenced global nuclear eviction upon laser-induced DSBs, and both the WT and mutant YAP2 showed similar translocation indicating that there may be additional factors beyond the Hippo pathway mediating this translocation. (Supp. Fig 3 A, B). We also directly assessed the levels of S127-phosphorylated YAP1 (pS127-YAP1) with IF. pS127-YAP1 was elevated in the nucleus following DNA damage, even as total YAP nuclear levels declined (Supp. Fig 3 C-E). Thus, even if YAP1 is phosphorylated at S127, somewhat surprisingly the phosphorylated form is not translocated under our experimental conditions. We also examined the phosphorylation of YAP at tyrosine 357 (Y357), which has been implicated in DNA damage associated YAP regulation and induction of apoptosis [33–36]. Following laser induced DNA damage, we do not observe any change in the nuclear to cytoplasmic ration of pYAP-Y357 within the time window corresponding to YAP eviction (Supp. Fig. 3 F, G), arguing against a role for Y357 phosphorylation in mediating nuclear export under these conditions. Since nuclear export represents one of the primary mechanisms by which proteins are transported from the nucleus, we asked whether YAP eviction on DNA damage might occur through an exportin1-dependent pathway. Exportin1 is well known to mediate the transport of leucine-rich nuclear export sequence (NES)-containing proteins, and inhibition of exportin1 by Leptomycin B has been shown to increase nuclear YAP levels under basal conditions [24, 44]. Using this inhibitor, we saw an expected increase in baseline YAP ratio, indicative of increased nuclear retention as expected. But causing DNA damage brought the YAP ratio down to similar to a similar extent in Leptomycin B treated cells when compared to control cells (Supp. Fig 3 H-J). Together these results point to an alternative mechanism for YAP nuclear eviction following DNA damage that operates independent of canonical Hippo signaling or active nuclear export pathways. Nuclear mechanics is emerging as a possible candidate for Hippo-independent control of YAP subcellular localization [20, 24] and this is what we investigated next.

**Figure 3.**
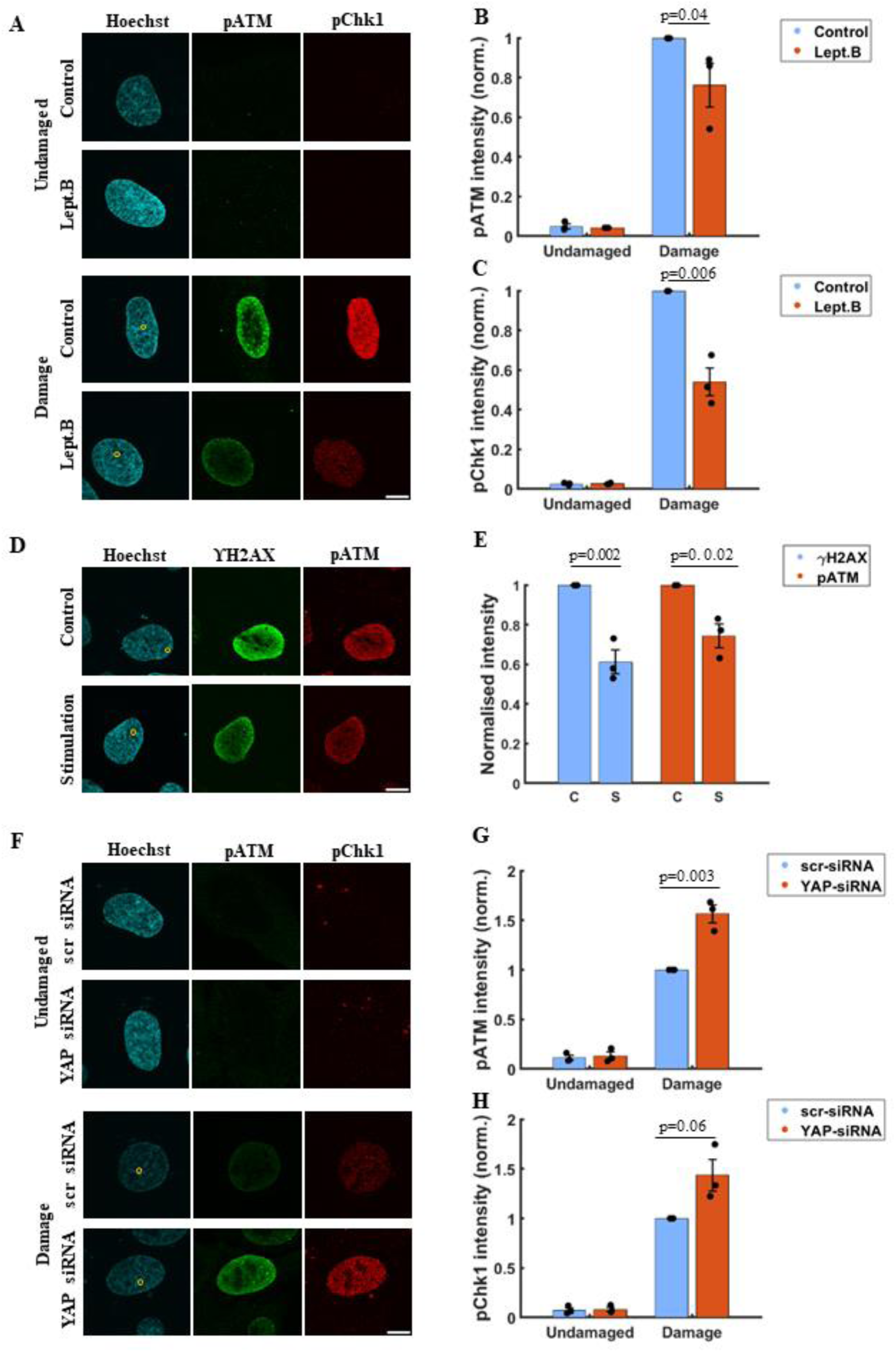
Nuclear YAP1 levels negatively correlate with DNA damage responses. (A) Representative images showing U2OS cell nuclei with Hoechst (cyan), intensities of endogenous pATM (pS1981) (green) and pChk1 (pS345) (red) in undamaged cells (top panel) and damage cells (bottom panel) of both vehicle treated (Control, Methanol:Water in Fluorobrite) and Leptomycin B treated (Lept.B) cells. The yellow circle represents the ROI of laser irradiation in the bottom panel. The scale bar is 10 microns. (B) Bar graph quantifying pATM levels of both vehicle treated (Control, blue) and Lept.B treated (Lept.B, orange) cells. Each dot represents the experiment-wise mean, normalised to the corresponding damaged control, and pooled across experiments (N=3, n>70 cells per condition). The p-values from the distributions are calculated using unpaired Student’s t-test. (C) Bar graph quantifying pChk1 levels in undamaged and damaged cells of both vehicle treated (Control, blue) and Lept.B treated (Lept.B, orange) cells. Each dot represents the experiment-wise mean, normalised to the corresponding damaged control, and pooled across experiments (N=3, n>70 cells per condition). The p-values from the distributions are calculated using unpaired Student’s t-test. (D) Representative images showing cell nuclei with Hoechst (cyan), intensities of endogenous ϒH2AX (green) and pATM (red) in OptoYAP overexpressing control and stimulated cells (Stimulation) on 405nm laser irradiation. The yellow circle represents the ROI of laser irradiation in the left panel. The scale bar is 10 microns. (E) Bar graph quantifying ϒH2AX levels (blue) and pATM levels (orange) in OptoYAP overexpressing control (C) and stimulated (S) cells on 405nm laser irradiation. Each dot represents the experiment-wise mean, normalised to the corresponding damaged control, and pooled across experiments (N=3, n>70 cells per condition). The p-values from the distributions are calculated using unpaired Student’s t-test. (F) Representative images showing U2OS cell nuclei with Hoechst (cyan), intensities of endogenous pATM (green) and pChk1 (red) in undamaged cells (top panel) and damage cells (bottom panel) in both scrambled siRNA (scr siRNA) and YAP siRNA treated (YAP siRNA) cells. The yellow circle represents the ROI of laser irradiation in the bottom panel. The scale bar is 10 microns. (G) Bar graph quantifying pATM levels in undamaged and damaged cells in both scrambled siRNA treated (scr siRNA, blue) and YAP siRNA treated (YAP siRNA, orange) cells. Each dot represents the experiment-wise mean, normalised to the corresponding damaged control, and pooled across experiments (N=3, n>70 cells per condition). The p-values from the distributions are calculated using unpaired Student’s t-test. (H) Bar graph quantifying pChk1 levels in undamaged and damaged cells in both scrambled siRNA treated (scr-siRNA) and YAP siRNA treated (YAP-siRNA) cells. Each dot represents the experiment-wise mean, normalised to the corresponding damaged control, and pooled across experiments (N=3, n>70 cells per condition). The p-values from the distributions are calculated using unpaired Student’s t-test.

### Nuclear mechanical changes on DNA damage affect YAP translocation and damage responses

YAP has been shown to be more nuclear under conditions of high nuclear tension [20]. Cytoplasmic YAP translocation thus can indicate relieving of nuclear prestress. The cell nucleus is held under a condition of prestress by the cytoskeleton [45, 46], and if the cytoskeletal tension is released one can expect effects even down to focal adhesions at the other end of cytoskeletal filaments [47]. We hypothesized that nuclear mechanics driven YAP relocalization might be a way to regulate YAP levels in the nucleus on DNA damage. If DNA damage causes mechanical changes, especially release of nuclear-cytoskeletal tension, one would expect in changes in the levels and arrangement of focal adhesion proteins. Indeed, we observed that the intensity of phospho-Focal Adhesion Kinase (pFAK), one of the major focal adhesion proteins, was lower in damaged cells compared to undamaged controls in the same plate (Fig. 2A, B) (Supp. Fig 4 A-C). We also observed a significant decrease in nuclear projection area over time in damaged cells compared to undamaged cells (Supp. Fig 4D) - this too is indicative of decreased nucleo-cytoskeletal tension. Next, we altered nuclear tension using an actin polymerization inhibitor Latrunculin A (LatA) to reduce actin-mediated prestress on the nucleus. LatA caused a significant decrease in YAP ratio, and causing further DNA damage in these cells resulted in no effect on YAP ratio, indicating that YAP ratio becomes insensitive to DNA damage on release of nuclear tension by F-Actin depolymerization (Fig, 2C, D). In addition, we enforced actin contractility by treated cells with Jasplakinolide, which stabilizes F-actin and prevents actin depolymerization, thereby increasing cytoskeletal and nuclear tension [48]. We observe a significant increase in YAP ratio in undamaged cells on Jasplakinolide treatment, consistent with increased mechanical loading of the nucleus. Importantly, upon laser micro-irradiation -induced DNA damage, cells treated with Jasplakinolide shows lesser loss of nuclear YAP compared to control (DMSO) treated cells (Supp. Fig 4 E, F). We next investigated what the effect of loss of nuclear YAP1 due to altered nuclear mechanics can be: is it just a byproduct of nuclear tension changes or does it have a functional role in effecting damage response? The ATM and ATR kinases are the main transducers of DSB response in cells [39]. On DNA damage, ATM is phosphorylated at S1981 [37] and ATR phosphorylates Chk1 [38] to initiate the cascade of DNA damage response. They can in turn, phosphorylate variant histone H2AX at Serine 139 (referred to as γH2AX) to amplify DNA damage signaling [49, 50]. To assess the impact of nuclear mechanical disruption, we examined γH2AX, phospho-ATM (pATM), and phospho-Chk1 (pChk1) levels in cells where nuclear mechanical forces were compromised. To specifically alter nuclear mechanical forces, rather than use a general cytoskeletal perturbant like LatA that can have other pleiotropic effects, we delinked the nucleoskeleton from the cytoskeleton using dominant negative Nesprin1 (dnNesprin1) and dominant negative LaminB1 (dnLaminB1) constructs. Nesprin proteins are positioned towards the outer nuclear membrane of the nucleus. Here, they bind to SUN domain proteins on the inner nuclear membrane and the cytoskeleton in the cytoplasm on the other [51, 52]. dnNesprin1 contains the C-terminal KASH domain from the Nesprin1 protein that binds to the SUN domain protein and lacks the actin binding domain. The binding of dnNesprin1 at the nuclear envelope saturates the available binding sites for endogenous Nesprin-1 by binding to SUN protein, resulting in their displacement from the nuclear envelope [18]. We also used a dominant negative LaminB1 construct, the overexpression of which is known to alter localization of Lamin A and C from the nuclear envelope, thus altering nuclear mechanical forces [53]. Overexpression of both constructs led to a decrease in baseline YAP ratio, though not to the extent that LatA does in undamaged cells. DNA damage under these conditions caused a further decrease in the YAP ratio (Fig. 2E, Supp. Fig. 4G). In dnNesprin1-overexpressing cells, DNA damage led to increased levels of γH2AX, pATM and a consistent but insignificant (p>0.05) increase in pChk1 (Fig. 2E-I). Similarly, dnLaminB1-overexpressing cells showed elevated γH2AX and pATM levels upon damage, whereas pChk1 remained unchanged (Fig. 2E-I). This suggests that nuclear mechanical forces help maintain YAP nuclear localization, but once these forces are sufficiently compromised, YAP is no longer retained in the nucleus, correlated with a difference in DDR kinase activity. The pan-nuclear γH2AX staining under these conditions could indicate subsequent apoptotic signaling [54], but such cell death if induced is expected at much later timepoints. To rule out if these mechanical changes upon DNA damage were not due to compromised cell membranes generally observed in dead/dying cells, we stained the cells with LIVE/DEAD cell imaging kit. This kit allows for simultaneous imaging of cell activity along with detection of dead/dying cells in different channels. 100% of the irradiated cells showed the marker for cell activity while 0% of the cells showed the cell death mark and they continue to show the mark for activity (Supp. Fig. 5 A,B). This indicates that the changes to nuclear mechanics observed are due to early DNA damage responses rather than long term processes of cell death, and the cells are viable and active at these timepoints. Together, these experiments suggest that nuclear mechanical changes can affect both the levels of YAP in the nuclei and DNA damage responses. But are these two readouts (YAP ratio and DDR activity) both merely correlated to nuclear mechanics, or is there a causative link between the two remained to be investigated.

**Figure 4.**
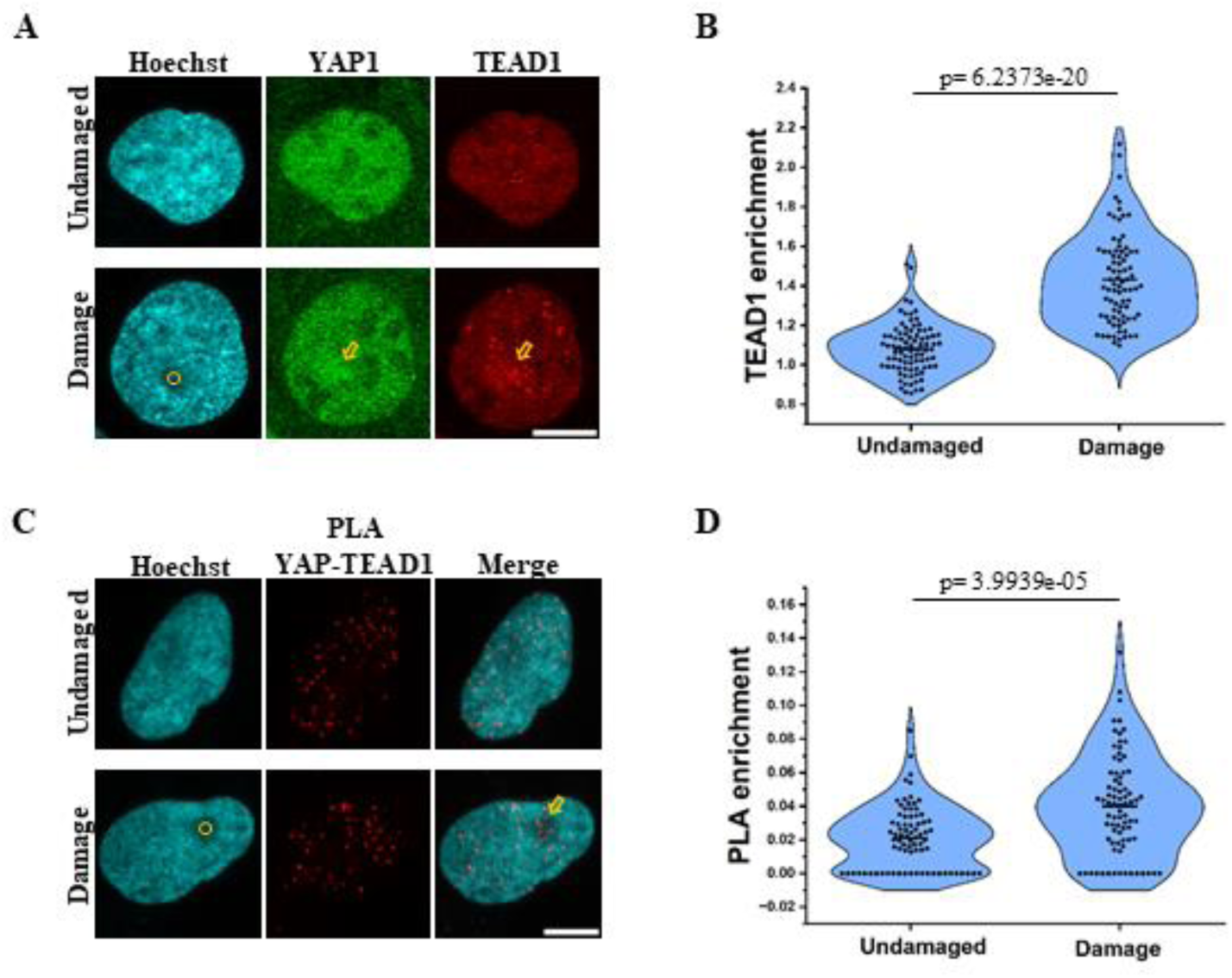
YAP selectively interacts with TEAD1 at sites of DNA Damage. (A) Representative images showing cell nuclei with Hoechst (cyan), intensities of endogenous YAP1 (green) and TEAD1 (red) in undamaged cells (top panel) and damage cells (bottom panel). The yellow circle represents the ROI of laser irradiation in the bottom panel. The yellow arrow indicates the enrichment of the proteins after damage. The scale bar is 10 microns. (B) Violin plot with distribution quantifying enrichment of TEAD1 intensity in undamaged and damaged cells. Each dot is the enrichment from a single cell. Enrichment is defined as the ratio of mean intensity at the site of damage to the mean intensity in the rest of the nucleus (N=3, n>70 cells per condition). For undamaged cell a random spot in the nucleus for each cell was chosen from the Hoechst channel as ROI. The p-values from the distributions are calculated using the Kolmogorov-Smirnov test. (C) Representative images showing cell nuclei with Hoechst (cyan), PLA intensity of YAP1-TEAD1 (red) and merge of the two channels in undamaged cells (top panel) and damaged cells (bottom panel). The yellow circle represents the ROI of laser irradiation in the bottom panel. The yellow arrow indicates the enrichment of the proteins after damage. The scale bar is 10 microns. (D) Violin plot with distribution quantifying enrichment of YAP1-TEAD1 PLA counts in undamaged and damaged cells. Each dot is the enrichment from a single cell (N=3, n>80 cells per condition). Enrichment is defined as the total number of PLA foci at the site of damage to the total number of PLA foci in the rest of the nucleus. For undamaged cell a random spot in the nucleus for each cell was chosen from the Hoechst channel as ROI. The p-values from the distributions are calculated using the Kolmogorov-Smirnov test.

**Figure 5.**
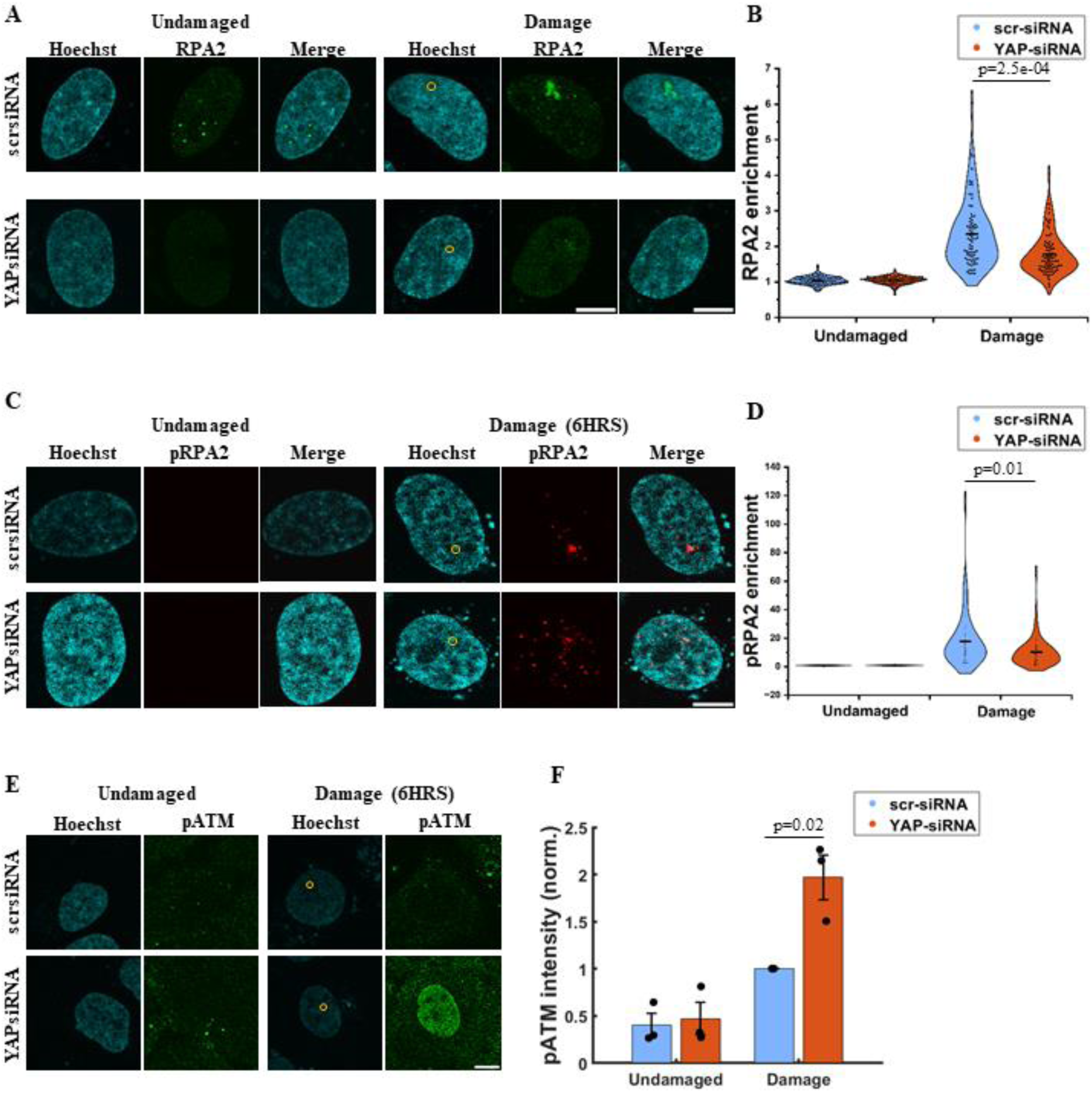
YAP depletion causes loss of DDR factors at site of damage and prolonged damage response. (A) Representative images showing U2OS cell nuclei with Hoechst (cyan), intensities of endogenous RPA2 (green) in undamaged cells (left panel) and damaged cells (right panel) in both scrambled siRNA treated (scr-siRNA, top) and YAP siRNA treated (YAP-siRNA, bottom) cells. The yellow circle represents the ROI of laser irradiation in the right panel. The scale bar is 10 microns. (B) Violin plot with distribution quantifying RPA2 enrichment in undamaged and damaged cells in both scrambled siRNA treated (scr-siRNA, blue) and YAP siRNA treated (YAP-siRNA, orange) cells. Each dot is the enrichment from a single cell (N=3, n>70 cells per condition). Enrichment is defined as the ratio of mean intensity at the site of damage to the mean intensity in the rest of the nucleus. For undamaged cell a random spot in the nucleus for each cell was chosen from the Hoechst channel as ROI. The p-values from the distributions are calculated using the Kolmogorov-Smirnov test. (C) Representative images showing U2OS cell nuclei with Hoechst (cyan), intensities of endogenous pRPA2(pS8) (red) in undamaged cells (left panel) and damage cells (right panel) in both scrambled siRNA treated (scr-siRNA, top) and YAP siRNA treated (YAP-siRNA, bottom) cells, 6 hrs after damage was caused. The yellow circle represents the ROI of laser irradiation in the bottom panel. The scale bar is 10 microns. (D) Violin plot with distribution quantifying pRPA2 enrichment in undamaged and damaged cells in both scrambled siRNA treated (scr-siRNA, blue) and YAP siRNA treated (YAP-siRNA, orange) cells. Each dot is the enrichment from a single cell (N=3, n>80 cells per condition). Enrichment is defined as the ratio of mean intensity at the site of damage to the mean intensity in the rest of the nucleus. For undamaged cell a random spot in the nucleus for each cell was chosen from the Hoechst channel as ROI. The p-values from the distributions are calculated using the Kolmogorov-Smirnov test. (E) Representative images showing U2OS cell nuclei with Hoechst (cyan), intensities of endogenous pATM (green) in undamaged cells (left panel) and damage cells (right panel) in both scrambled siRNA treated (scr-siRNA, top) and YAP siRNA treated (YAP-siRNA, bottom) cells, 6 hrs after damage was caused. The yellow circle represents the ROI of laser irradiation in the right panel. The scale bar is 10 microns. (F) Bar graph quantifying pATM levels in undamaged and damaged cells in both scrambled siRNA treated (scr-siRNA, blue) and YAP siRNA treated (YAP-siRNA, orange) cells. Each dot represents the experiment-wise mean, normalised to the corresponding damaged control, and pooled across experiments (N=3, n>70 cells per condition). The p-values from the distributions are calculated using unpaired Student’s t-test.

### Nuclear YAP1 levels negatively correlates with DNA damage responses

To understand the specific role of YAP in DDR, we modulated the levels of YAP in the nuclei and observed the effect on DNA damage responses. First, we increased the levels of YAP in the nuclei using Leptomycin B. As discussed earlier, Leptomycin B treatment leads to increase in nuclear YAP levels. Correspondingly, under such treatment conditions, both pATM and pChk1 levels in the nuclei are lower compared to vehicle-treated cells upon DNA damage (Fig. 3A-C). Leptomycin B can influence other cellular processes beyond promoting YAP nuclear accumulation, which could indirectly affect the DNA damage responses. To rule out this possibility, and specifically modulate YAP levels, we employed an optogenetic system (OptoYAP) that allows precise control of YAP nuclear localization in a light-dependent manner [55]. OptoYAP construct consists of the LOV2 and Jα domains fused to YAP-mCherry. In the basal state, these domains interact such that a nuclear localization signal (NLS) is masked while a nuclear export signal (NES) remains exposed, thereby restricting YAP to the cytoplasm. Upon 488 nm light stimulation, LOV2-Jα domains unfold, unmasking the NLS and promoting YAP nuclear import, which can be visualized via the mCherry tag (Supp. Fig. 6 A-C). We stimulated cells to cause increased YAP nuclear localization, subsequently irradiated using a 405nm laser to cause DNA damage and the levels of ϒH2AX and pATM were observed after 20 minutes. As a control, cells which were not stimulated but exposed to 405nm laser to cause damage were used. As expected, 488 nm stimulation led to a robust increase in nuclear YAP accumulation, whereas control cells showed no change (Supp. Fig. 6 B, C). Notably, cells with optogenetically driven nuclear YAP (Stimulated, S) displayed significantly reduced ϒH2AX and pATM levels compared to unstimulated control (C) (Fig. 3D, E), in line with the studies using Leptomycin B, arguing for a role of YAP in DDR. To further establish the relationship between YAP and DDR, we decreased the levels of YAP in the cell by using siRNA against YAP (Supp. Fig 7 A, B). We first verified if YAP siRNA could alter cell cycle composition and potentially influence DDR signaling. To address this, we performed flow cytometry-based cell-cycle profiling of cells transfected with scrambled (scr) siRNA and YAP siRNA under identical conditions. DNA content analysis revealed no significant differences in the distribution of cells across G1, S and G2/M phases between scr transfected and YAP depleted populations (Supp. Fig. 7A). Under such YAP knockdown conditions, DNA damage caused increased ϒH2AX, pATM and a consistent but insignificant (p>0.05) increase pChk1 intensity in the nucleus, compared to scrambled siRNA treated cells, with the increasing trend observed upon increasing doses of DNA damage (Fig. 3F-H; Supp. Fig 7 C, D), in line with the changes observed with increasing nuclear YAP. In addition to using the exact same irradiation protocol we also verified that the extent of primary damage in scrambled and YAP siRNA treated cells were similar. For this we performed Terminal Deoxynucleotidyl Transferase dUTP Nick End Labelling (TUNEL) assay to assess for the extent of primary damage i.e. strand breaks. The TUNEL protocol stains for the 3’-OH termini generated during single and double strand breaks. Using TUNEL assay, we could visualize the strand breaks at the site of irradiation, and did not see any difference in the extent of DNA damage between scrambled and YAP siRNA treated cells, as measured by the enrichment of TMR RED intensity at the site of damage (Supp. Fig. 7 E, F). Thus, increased nuclear YAP corresponds to lower DDR activity and vice versa. We also wondered if these observations are specific to laser-induced clustered DSBs or can they be generalized. To address this, we treated cells with Neocarzinostatin, a DNA-damaging agent that induces DSBs. Even under this damage condition, YAP knockdown cells exhibited elevated ϒH2AX levels compared to control scrambled siRNA-treated cells (Supp. Fig. 7 G, H). These results collectively suggest that nuclear YAP levels are critical modulators of the DNA damage response, with lower nuclear YAP levels corresponding to increased DNA damage signaling by regulating the activity of critical DDR kinases.

**Figure 6.**
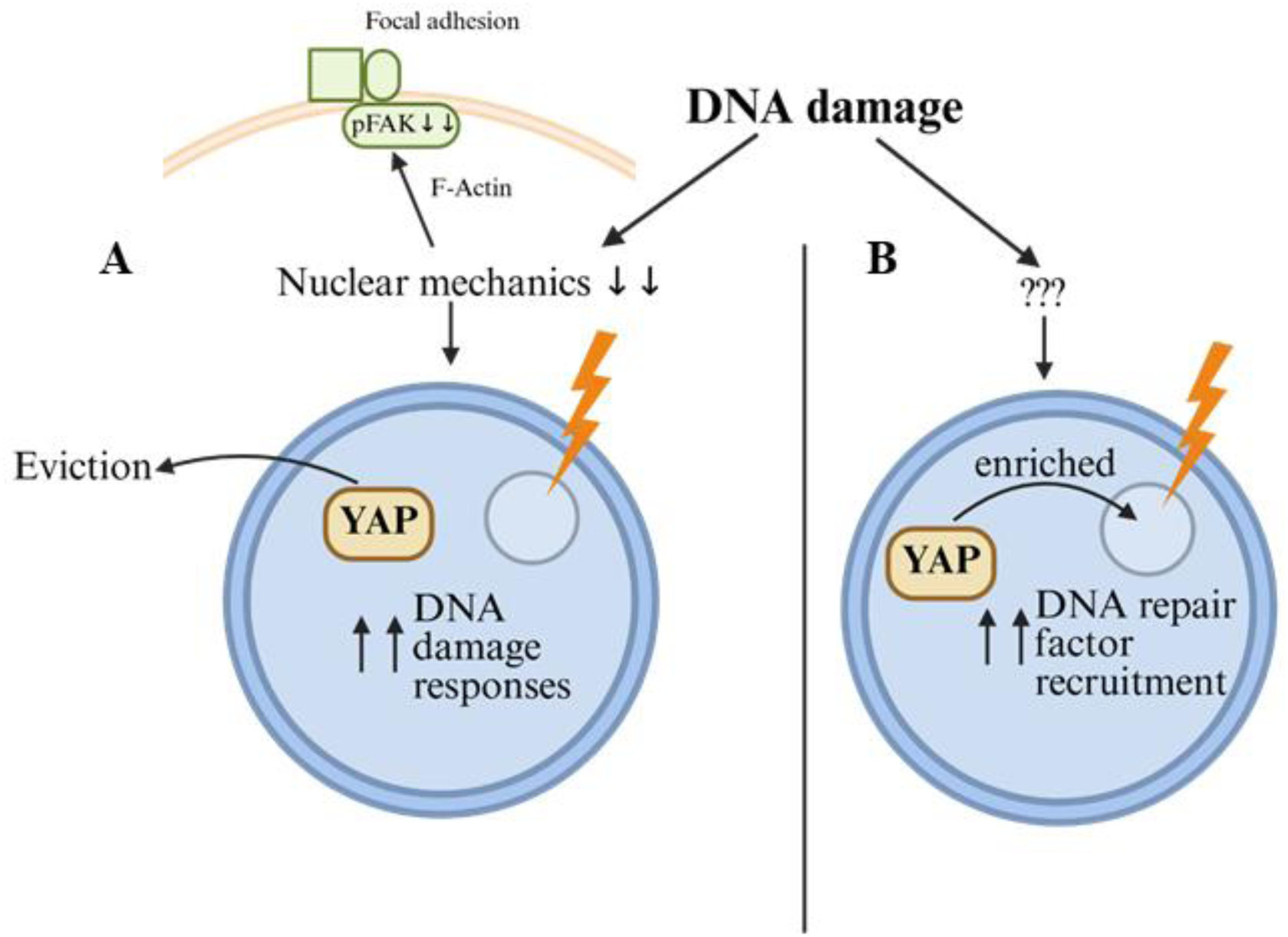
Proposed model for role of YAP in regulating DNA damage responses. (A) Nuclear mechanical forces are decreased following DNA damage, resulting in eviction of YAP from the nucleus, which enhances DNA damage responses. (B) DNA damage parallelly also leads to the enrichment of YAP at DNA lesions, which promotes the recruitment of DNA repair factors.

### YAP Selectively Interacts with TEAD1, but not ARID1A or p73, at Sites of DNA Damage

While we described global eviction of YAP under laser-induced damage, intriguingly we had also observed YAP to be locally enriched at the site of DNA damage (Fig 1E, G) (Supp. Fig 2 A, C). We wondered if this global reduction and local enrichment serve a different purposes in modulating DDR. We thus next looked for known interactors of YAP that are known to bind DNA repair factors and whether they interact with YAP on DNA damage. ARID1A (part of the SWI/SNF complex), TEAD1 and p73 (transcription factors known to associate with YAP to promote proliferative and apoptotic programs respectively) under different conditions have been shown to interact with YAP in independent studies [25, 56, 57] to either bring about chromatin remodeling (in the case of ARID1A), increase in proliferation (in the case of TEAD1) or signal apoptosis (in the case of p73) [58, 59]. We first looked at their site enrichment at sites of DNA damage. We did not observe any induction of p73 on DNA damage under our conditions (Supp. Fig 8 A, B). To confirm antibody functionality, we treated cells with Etoposide under conditions previously shown to induce p73 expression [58, 59]. We observed a robust induction of the antibody under such conditions (Supp. Fig 8 C, D). In contrast to p73, we observed that both ARID1A and TEAD1 are recruited to the site of DNA damage (Fig 4 A, B) (Supp. Fig 9 A, B). To probe for interaction between YAP-ARID1A and YAP-TEAD1, we performed Proximity Ligation Assay (PLA). This assay allows for the visual detection of protein-protein interactions. Antibodies specific to the proteins of interest are conjugated with short DNA oligonucleotides, which can then be ligated, amplified and fluorescently probed to detect interaction. We investigated both site specific and global interaction of YAP-TEAD1 and YAP-ARID1A on DNA damage using PLA. We find that at 20 min time point after damage, there was a significant increase in the local enrichment of YAP-TEAD1 (Fig 4 C, D), while no significant change was observed for YAP-ARID1A interaction on DNA damage (Supp. Fig 9 C, D). The overall nuclear counts of PLA remained constant in either case (Supp. Fig 9 E, F). This shows that YAP enriches and interacts with TEAD1 at the site of DNA damage, and because TEAD1 has been shown interact with DNA repair factors [59], this interaction may have a direct role in DDR at the site. This implies that knocking down YAP1 may influence repair factor recruitment at the damage site, beyond its global effect on DDR kinase activity, which is what we investigated next.

### YAP depletion causes loss of DDR factors at site of damage and persistent damage response

Given the results above, we aimed to investigate the consequences of YAP recruitment to the site of DNA damage. The first step was to identify candidate DNA damage response proteins that can potentially directly bind to YAP. Recently, Chang et. al. had identified several nuclear proteins that can interact with YAP by performing YAP immunoprecipitation followed by mass spectrometry [25]. We sorted out DNA damage response proteins from that list (Supp. Fig. 10A). These candidate proteins that interact with YAP could be brought in by YAP at the site of DNA damage, thus helping in DNA damage repair. We then performed immunofluorescence after DNA damage to observe the site recruitment of these proteins in control cells or when YAP levels were depleted with YAP siRNA. Among the proteins we investigated, we observed the levels of RPA2 to be significantly reduced at the site of DNA damage on YAP knockdown (Fig. 5 A, B). To check if this reduction persists at longer time points, we stained for both RPA2 and the phosphorylated form of RPA2 involved in DNA damage repair (pS8-RPA2) [60–63] at 6hrs after damage. We observed that the decreased site recruitment of RPA2 and pRPA2 in YAP siRNA cells persisted for a longer time after DNA damage (Fig. 5 C, D; Supp. Fig. 10B). Finally, to understand the long-term consequences of DNA damage response in the absence of YAP, we stained for pATM 6hrs after DNA damage. We observed persistence of pATM intensity in cells treated with YAP siRNA (Fig. 5 E, F). This indicates that global DDR signaling persists longer while the accumulation of local repair factors is reduced in the absence of YAP.

## Discussion

Our study suggests two ways by which YAP can affect DNA damage responses. (1) Eviction of YAP from the nucleus through changes in nuclear mechanics causes an increase in ATM and ATR activity. Thus, while DDR and YAP-dependent programs are independently known to connect to nuclear mechanics, they may also be interconnected by the common thread of mechanics (2) The remaining fraction of YAP that is still nuclear, enriches at the site of DNA damage facilitating repair by recruiting DNA repair factors. Thus together, the global eviction and local enrichment could both ensure efficient repair of the damaged DNA and effective DDR (Figure 6).

Our experiments suggest a novel way in which DNA damage induced changes in nuclear mechanics can regulate DNA damage response by controlling the nuclear YAP levels. This eviction was consistently observed for both isoforms of YAP, YAP1 and YAP2. Nuclear eviction of YAP starts within seconds of DNA damage (Fig. 1C), suggesting us that it might be regulated by mechanical forces rather than slower biochemical signaling. While we saw an increase in phosphorylation of YAP at S127 at earlier time points, it does not appear to be the main driver of its early nuclear eviction as it is itself nucleus-confined. It has been shown that DNA damage causes association between LATS and YAP to trigger changes in binding ability of YAP with its transcription factors [30]. Notably, the fraction of YAP that is evicted from the nucleus is not phosphorylated at S127, as shown by the similar nuclear decrease of wild-type YAP2 and S127A mutant YAP2 upon DNA damage. Exportin-1 does not appear to mediate YAP nuclear export on DNA damage either. Leptomycin B treatment causes increase in nuclear YAP levels in line with exportin1 inhibition, but the eviction upon DNA damage still occurs. Our study provides a strong argument for changes in nuclear mechanics to be a key regulator of YAP export from the nucleus upon damage. We hypothesize that alongside Hippo signaling, YAP may be evicted via changes in nuclear pore permeability [24], with decreased cytoskeletal tension upon DNA damage as suggested before.

Changes in nuclear mechanics using Atomic Force Microscopy (AFM) upon DNA damage by long time treatment with cisplatin has been reported recently [26]; however, the mechanism by which it affects DNA repair remain unclear. We demonstrate that global eviction of YAP via altered nuclear mechanics enhances DNA damage responses, as seen by elevated γH2AX, pATM and pChk1 levels with both laser-induced and chemically induced DSBs. Nuclear mechanics is emerging as a key regulator of cellular processes like chromosome movement, differentiation, development, cell competition [2, 64–67]. Emerging strategies that aim to soften the tissue matrix and thereby the nucleus, are being explored to limit metastatic progression and cancer development [68]. Our findings suggest that decrease in nuclear mechanics and resultant YAP eviction increases DNA damage responses. The nucleus is mechanically coupled to the cytoskeleton through structural links in place like the LINC complex, allowing changes in cellular or substrate rigidity to influence nuclear morphology and stiffness [18]. Among cytoskeletal filaments, F-actin plays a major role in regulating nuclear mechanics. We show that DNA damage causes changes in pFAK levels at focal adhesions, indicative of loss of actin-mediated nuclear tension. Our data also show that F-actin-mediated forces help retain YAP in the nucleus following damage. Upon Latrunculin A mediated disruption of F-actin, the nucleus loses its ability to retain YAP in the nucleus and YAP ratio is not affected further on DNA damage, reaching a global minimum of low mechanics. On specifically altering nuclear mechanical forces using dnNesp1-mApple and dnLamB1-mApple constructs, we found that YAP levels were already significantly reduced even without DNA damage and continued to decrease upon damage. dnLaminB1 had a more pronounced effect than dnNesprin1 on YAP1 levels in control cells as well its subsequent damage-induced decrease, possibly due to its ability to disrupt both Lamin A and C. Regardless, mechanical disruption by either construct elevated DDR.

We confirmed that YAP is a critical mediator of nuclear mechanotransduction during DNA damage response. Changing YAP levels both ways (increase or decrease) influenced damage signaling: nuclear YAP increase on Leptomycin B treatment or through optogenetic means, suppressed ATM/ATR activity, whereas siRNA-mediated knockdown of YAP increased it. This increase is apparent even at lower DNA damage, as shown by the trend of increasing γH2AX intensity on YAP knockdown (Supp. Fig. 7 C). This indicates that YAP may regulate critical DDR kinases, possibly by controlling their upstream regulators. Therefore, nuclear eviction of YAP appears to be a regulatory mechanism for activating the DNA damage response globally. Apart from interactions with DDR factors, YAP of course can have myriad transcriptional targets depending on the partner recruiting it to DNA - for example TEAD1 can bring about a proliferative program, while p73 can bring about a pro-apoptotic program. Global YAP eviction can thus potentially reduce pro-proliferative programs, while giving cells the opportunity to repair damage or die by apoptosis. We do not see signatures of cellular apoptosis, as measured by p73 intensity (Supp. Fig 8A, B) or in a LIVE/DEAD assay (Supp. Fig 5), and indeed the cells are alive in the time course of our experiments.

Beyond global nuclear eviction, YAP is also enriched at DNA damage sites across varying levels of damage. Although the mechanism of this recruitment in unknown, it is functionally relevant. While eviction of YAP occurs significantly at higher extent of DNA damage, its local enrichment occurs both at lower and higher dose of DNA damage in a dose dependent manner (Supp. Fig 2 A, C). Both TEAD1 and ARID1A are recruited with YAP to varying degrees at the site of damage, and YAP interacts with TEAD1 as shown by proximity ligation assay, potentially establishing a favorable environment at DNA damage site to facilitate repair. TEAD1 has recently been shown to be important in DNA repair and its depletion decreases DNA repair efficiency [59]. We show that TEAD1 is indeed recruited to damage sites and may work with YAP to enhance repair through multi-protein assemblies. We did not observe YAP enrichment at the damage site using EGFP-YAP1 overexpression constructs, possibly due to interference from the EGFP tag. This issue could potentially be overcome in future experiments using anti-YAP nanobodies as live cell tracers of YAP.

Using a prior study, we identified eight DNA repair proteins which interact with YAP [25] (Supp. Fig 10A). Of the ones we investigated, RPA2 enrichment was most affected by YAP knockdown. RPA2 and pRPA2 recruitment was impaired at early and late time points respectively following damage, suggesting impaired repair. RPA2 binds single stranded DNA generated when Double Strand Breaks (DSBs) are repaired. In addition to protecting the exposed DNA to nucleolytic cleavage, it has also been shown to direct damage response towards Homologous Recombination (HR) repair [69]. Phosphorylation at S8 of RPA2 has been shown to be important in DNA repair, replicon stability and checkpoint progression [60–63]. Thus, the presence of YAP might dictate the repair outcome by bringing in DNA repair factors. Additionally, elevated pATM levels persisted even six hours after damage in YAP knockdown cells. These findings suggest that YAP plays dual roles on DNA damage-globally promoting damage signaling via its nuclear absence and locally promoting repair at the damage site. This duality not only deepens our understanding of YAP function beyond its canonical roles but also highlights potential opportunities to modulate DNA repair responses in diseases such as cancer, where mechanical environments and DNA repair capacity are often altered. Together our study highlights a broader involvement of YAP in maintaining genomic stability than previously appreciated.

YAP/TAZ are increasingly being recognized as important regulators of tumor initiation, progression and metastasis [3]. Although YAP/TAZ activity is often dispensable for normal tissue homeostasis, many cancers exhibit a strong dependence on YAP/TAZ signaling for proliferation and survival [70]. Mechanistically, YAP/TAZ regulate transcriptional programs controlling cell cycle progression, DNA synthesis, mitotic completion and they can cooperate with proto-oncogenic transcription factors like c-Myc to reinforce malignant phenotypes [71]. In addition, YAP/TAZ activity has been linked to resistance to anoikis through regulation of Bcl2 family members, as well as to reduced sensitivity to chemotherapy and radiotherapy [72]. Notably, dysregulated YAP/TAZ signaling is frequently observed in human tumors even in the absence of canonical Hippo pathway mutations [73], indicating that alternative regulatory inputs, including mechanical cues, might play a major role in controlling YAP activity in cancer.

Our findings suggest an additional dimension to YAP function in tumor biology by linking nuclear mechanics, YAP localization and DNA damage responses. We find that global nuclear eviction of YAP enhances DDR signaling, whereas site specific recruitment of YAP prolongs damage signaling. These observations raise the possibility that aberrant YAP regulation in tumors with elevated nuclear YAP, commonly observed in gliomas, breast and colorectal cancers, may alter the balance between damage signaling and repair, potentially contributing to genomic instability or therapy resistance. From a translational perspective, these findings suggest that targeting the YAP-DDR and YAP-TEAD1 axis may represent a strategy to modulate tumor sensitivity to genotoxic therapies. Our results further suggest that mechanical regulation of YAP, for example through pathways controlling nucleo-cytoskeletal coupling, could provide an additional point of therapeutic intervention. Modulating nuclear mechanics of YAP localization may therefore represent a means to enhance the efficacy of radiotherapy or DNA damaging agents in tumors characterized by high YAP activity.

## Materials and methods

### Cell culture

U2OS cells were obtained from ATCC (via HiMedia Laboratories) and cultured in McCoy 5A media (Sigma, M4892) supplemented with 10% FBS (Gibco, 16000-044), 1% PenStrep-glutamine (Gibco, 10378-016) and sodium bicarbonate. Cells were maintained in T-25 flasks (Tarsons, 950040) and incubated at 37°C and 5% CO2 in a CO2 incubator (Eppendorf Galaxy 170S). For experiments and imaging, the cells were plated in a glass-bottom dish (Cellvis) at least 24 hours prior to experiments to achieve moderate confluency. For live cell experiments, cells were imaged in FluoroBrite medium (Gibco, A1896701) supplemented with 25mM HEPES, 10% FBS (Gibco, 16000-044) and 1% PenStrep-glutamine (Gibco, 10378-016). Plates were placed in the live-cell chamber and maintained at 37°C. Transfections were performed using XtremeGENE™ HP DNA Transfection Reagent (Roche, 6366236001) following the manufacturer’s protocol.

### Immunofluorescence

Cells were grown as described previously. Following damage, cells were allowed to recover for 20mins unless mentioned otherwise. Cells were then fixed using 4% PFA (paraformaldehyde, Sigma P6148) in 1X PBS (phosphate-buffered saline, Sigma P5493) for 10 minutes. PFA was removed, and cells were washed twice with 1X PBS. Permeabilization was carried out using 0.3% Triton X-100 (Sigma, T8787) in 1X PBS for 15 minutes, followed by two washes with 1X PBS. Cells were then blocked with 5% BSA (bovine serum albumin, Himedia TC545) in 1X PBS (Blocking solution) for an hour. Primary antibodies (with dilution as mentioned in the table) were diluted in the blocking solution and incubated overnight at 4°C. The following day, primary antibodies were removed and cells were washed twice with 1X PBS. Secondary antibody incubation was subsequently done in the blocking solution for 1 hour at room temperature. Finally, cells were washed twice and re-imaged in 1X PBS for damaged cell fluorescence intensities along with an equal number of undamaged cells in the same plate.

### Cell treatments

The following inhibitors and their working concentrations are as follows: Leptomycin B (L2913-Sigma) at 20nM for 4hrs, Latrunculin A (L5163-Sigma) at 0.5 μM for 30mins, Jasplakinolide (J7473-ThermoFisher) at 150nM for 40mins. The inhibitors were treated for the stated time and imaging was performed in the presence of the inhibitors to prevent recovery. To induce DNA damage and assess YAP nuclear-to-cytoplasmic ratio changes, Neocarzinostatin (NCS) was used at a concentration of 10 ng/mL for 4 h. For siRNA experiment NCS was used at a working concentration of 1μg/ml for 1hr in media. Etoposide was used at a working concentration of 100μM for 24 hours. Staurosporine was used at a working concentration of 1μM for 4 hours and then stained for LIVE/DEAD stain.

### TUNEL assay

The assay was performed using the In Situ Cell Death Detection Kit, TMR red (Roche, Cat. No. 12156792910). This assay helps to detect and quantify DNA strand breaks at single-cell level by labelling the free 3’-OH termini of strand breaks with modified nucleotides using the enzyme Terminal deoxynucleotidyl transferase followed by detection using fluorescent imaging. Fluorescence intensity and enrichment were quantified on a cell-by-cell basis at the stated time points after damage to assess damage persistence and repair.

### Flow cytometric analysis

Cells were grown on a 35 mm plastic-bottom plate till confluency. After which the cells were detached from the surface with trypsin. The cells were then transferred into a 15 ml falcon tube and trypsin was de-activated by Full Medium (DMEM (Dulbecco Modified Eagle Medium) with 10% FBS (fetal bovine serum), 1% penicillin and streptomycin) added thrice the quantity of trypsin used for the detachment of the cells. The cell mix was then spun at 300g for 10 minutes and supernatant was discarded. The pellet was resuspended in 4% PFA (para-formaldehyde) in PBS (phosphate buffered saline) for fixation (10 minutes). The cell-PFA mix was again spun at 300xg for 10 minutes and supernatant was discarded. The pellet then was resuspended in 0.3% Triton X-100 in PBS (10 minutes) for permeabilization. It was followed by spinning at 300xg for 10 minutes. The supernatant was discarded and the pellet was resuspended with 2 *μ*g/ml DAPI in PBS. After incubation of 7 - 10 minutes with DAPI, the cell-DAPI mix was spun at 300g for 10 minutes and the pellet obtained was resuspended in PBS and was taken to flow cytometer (Cytoflex S, Beckman Coulter) and analyzed using 405nm laser excitation. Analysis was done using FloJo 10 software.

### LIVE/DEAD cell imaging kit

Cells were cultured in dishes as described previously. Following damage and 20mins incubation, the Live Green vial (A) and the Dead Red Vial (B) from the LIVE/DEAD cell imaging kit were mixed to create a 2X stock, following which it was mixed with equal volume of Fluorobrite and incubated in the dish for 15mins in tissue culture hood at room temperature. The dish was then re-imaged for the damaged cells and an equal number of undamaged cells, within 1 hr of treatment for intensity of LIVE stain in the FITC channel and DEAD stain in the TexasRed channel.

### Confocal microscopy and micro-irradiation

Imaging was performed on a confocal microscope (Olympus FV3000) unless mentioned otherwise. Double Strand Breaks (DSBs) were induced in Hoechst-sensitized cells with 405 nm laser as described before [74, 75]. Fluorescence images were acquired using a 60× oil objective (PlanApo N 60× Oil, numerical aperture = 1.42, Olympus) with a 4.5× optical zoom (except where stated otherwise) mounted on an Olympus IX83 inverted microscope equipped with a scanning laser confocal head. Cells grown in 35-mm glass-bottomed dishes were sensitized with 1µg/ml Hoechst for 10 minutes in media, washed off and subsequently supplemented in FluoroBrite for imaging. The plate was positioned on the microscope’s live cell chamber for 15 minutes prior to imaging. A point region of interest (ROI) was placed on a cell-by-cell basis and DSBs were induced with the 405 nm laser. Post fixing and staining, coordinates of cells were re-loaded and the most focused frame was captured for the mentioned antibodies in 512 × 512 resolution. The laser power for 405nm laser measured at 60x objective at 100% power was 0.874mW. Laser irradiation was performed at 20% intensity of the laser (dwell time 250ms, speed 4μs/pixel), unless mentioned otherwise. For pFAK image acquisition, stained cells were imaged along the z-axis (0.5μm steps - 9 steps), along with equal number of undamaged cells from the same plate.

### OptoYAP experiments

Hoechst-sensitized U2OS cells overexpressing OptoYAP (pDN34-BiNLS-hYAP1-1delta) were imaged using a 561 nm laser to record baseline localization. Following this, half of the cells were subjected to optogenetic stimulation with a 488 nm laser applied every 1 min for 30 min over a rectangular ROI encompassing the entire cell. After stimulation, all cells were re-imaged with the 561 nm laser to assess OptoYAP nuclear enrichment. Subsequently, localised DNA damage was induced in all cells using a 405 nm laser at 20% intensity, applied to a point ROI, followed by incubation for 20 min in a live-cell chamber. Cells were then fixed and immunostained with the indicated antibodies.

### In-situ proximity ligation assay (PLA)

PLA was performed using Duolink In-situ reagents (Sigma-Aldrich). After microirradiation of Hoechst sensitized U2OS cells plated on 35mm glass bottom dish, cells were fixed using 4% PFA in 1X PBS for 10 minutes and washed twice with 1X PBS. Permeabilization was performed using 0.3% Triton X-100 in 1X PBS for 15 minutes followed by two 1X PBS washes. Subsequent steps were carried out using the manufacturer’s instructions. Coordinates of damaged cells were reloaded and imaged along the z-axis (0.5μm steps - 11 steps), along with equal number of undamaged cells from the same plate. Image stacks were average-projected prior to analysis.

### RNA interference

YAP knockdown experiments were performed using an ON-TARGETplus SMARTpool siRNA from Dharmacon (Ref #: SO-3203873G). It consists of four independent siRNAs targeting YAP. ON-TARGETplus reagents are specifically designed to minimize off-target effects through chemical modifications that reduce seed-sequence-mediated micro-RNA like interactions and through sequence selection algorithms that avoid transcripts with high off-target potential. In addition, the use of a pooled format allows gene silencing to be achieved at lower concentrations of each individual siRNA, which further reduces nonspecific effects while maintaining robust knockdown efficiency [76, 77]. siRNA transfections were done using X-tremeGENE siRNA transfection reagent (Sigma-Aldrich) in Opti-MEM, following manufacturer’s protocols. YAP siRNA was used at 300nM final concentration. Media was replaced 2 days after transfection and experiments were performed 3 days post media change. Cell cycle composition was comparable between the scrambled and YAP siRNA transfection populations (Supp. Fig. 7 A).

The sequences of the siRNAs used in the SMARTpool are:

a) GCACCUAUCACUCUCGAGA
b) UGAGAACAAUGACGACCAA
c) GGUCAGAGAUACUUCUUAA
d) CCACCAAGCUAGAUAAAGA

### Image analysis

Multichannel images were separated using Fiji. Cell-by-cell intensity quantification was done using an in-house MATLAB (Mathworks) script. For figure representation, image contrast was uniformly adjusted in Fiji across all experimental groups. Nuclear-to-cytoplasmic (N/C) ratios of YAP were calculated as follows: a binary nuclear mask was generated from the Hoechst channel and multiplied with the YAP channel to obtain nuclear intensity. To define the cytoplasmic region, the nuclear mask was dilated, and the inverted undilated mask was multiplied to exclude the nucleus, yielding a cytoplasmic mask of constant area surrounding the nucleus. Cytoplasmic intensity was then measured from this mask, and the nuc./cyto. ratio was calculated as nuclear intensity divided by cytoplasmic intensity. For pFAK image quantification, z-stacks were average projected, and a binary mask of focal adhesions was generated from the pFAK channel. The binary mask was then multiplied to the original image get the total intensity at the focal adhesions. For time-lapse analysis of cells imaged in the absence of Hoechst staining, a nuclear channel was not available. Therefore, nuclei were manually segmented at each time point using the YAP channel, which allowed identification of the nuclear region based on the characteristic nuclear enrichment of YAP signal. This approach enabled consistent tracking and quantification of nuclear YAP intensity in the no-Hoechst control condition across the time series. The same segmentation and analysis strategy was also applied to the Hoechst-stained samples to ensure direct comparability between Hoechst and non-Hoechst conditions.

### Statistical analysis

All statistical analyses were performed in MATLAB. Bar and line plots were generated using MATLAB, while violin plots were generated using Origin. Bar graphs represent the mean along with standard error of the mean (SEM) unless mentioned otherwise. Comparison of means from multiple experiments was done using a Student’s t-test. Intensity normalizations are done to the respective damaged controls, as all the undamaged samples have very low levels of staining making normalization to those difficult. Single cell data often may not conform to the assumption of normality, thus for single cell data, non-parametric test (Kolmogorov Smirnov’s test) was employed. Although individual cells are represented as data points in the violin plots, the observed trends were first verified to be consistent across biological replicates before pooling single-cell measurements for statistical analysis. Single cell data was pooled only for ratios (nuclear to cytoplasmic ratios or enrichment scores) that are independent of baseline staining. Kolmogorov-Smirnov tests were used for comparing single cell level data as such non-parametric tests do not make assumptions of normality about the underlying distributions. On the other hand, when means from multiple repeats of an experiment are compared (say for staining intensities), Student’s t-tests were used. Here the data satisfy the assumptions of the test. Statistical tests used for comparisons are indicated in the figure legends. In all our analyses, “N” refers to the number of independent biological replicates, while “n” refers to the total number of individual cells analyzed per condition in those replicates.

### Antibodies and plasmids

The following plasmids were used in this study: pEGFP-C3-hYAP1 (Addgene plasmid # 17843), pDsRed Monomer C1-YAP2 (Addgene plasmid # 19057) and pDsRed Monomer C1-YAP2-S127A mutant (Addgene plasmid # 19058) was a gift from Marius Sudol. OptoYAP (pDN34-BiNLS-hYAP1-1delta) was a gift from Timothy Saunders (Addgene plasmid # 169471; http://n2t.net/addgene:169471; RRID: Addgene_169471). mApple-dnNesprin1 (amino acids 8369-8749 of human Nesprin-1 cloned into mApple-C1) (DOI:10.1242/jcs.02471), mApple-dnLaminB1 or XLaminB1Δ2+ (34-420 aa from X06344 cloned into mApple-C1) (PMID: 11683386) and C1-mApple (Addgene #54631) was a gift from Tamal Das. GFP-Nucleolin was a gift from Michael Kastan (Addgene plasmid # 28176; http://n2t.net/addgene:28176; RRID: Addgene_28176).

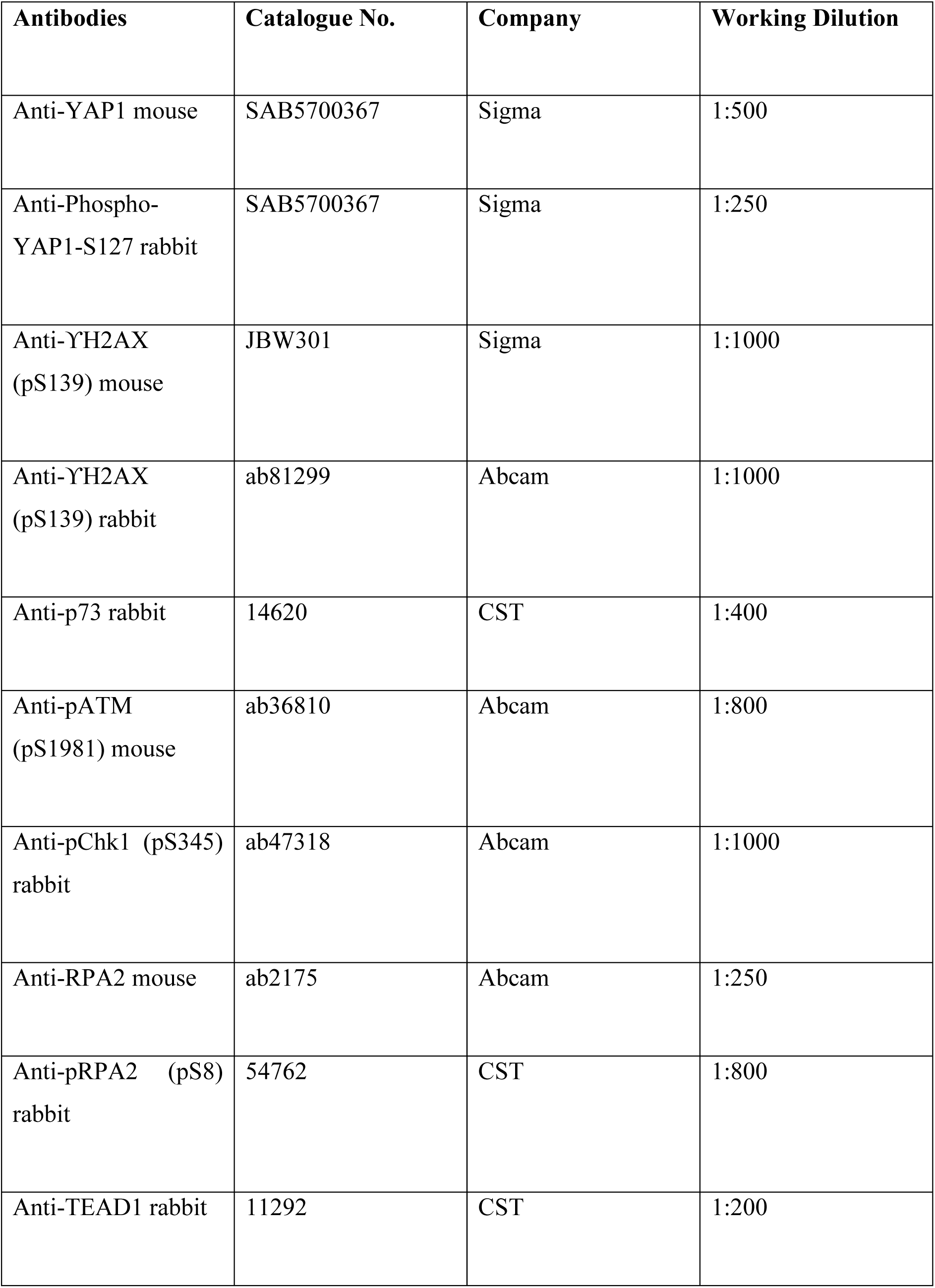
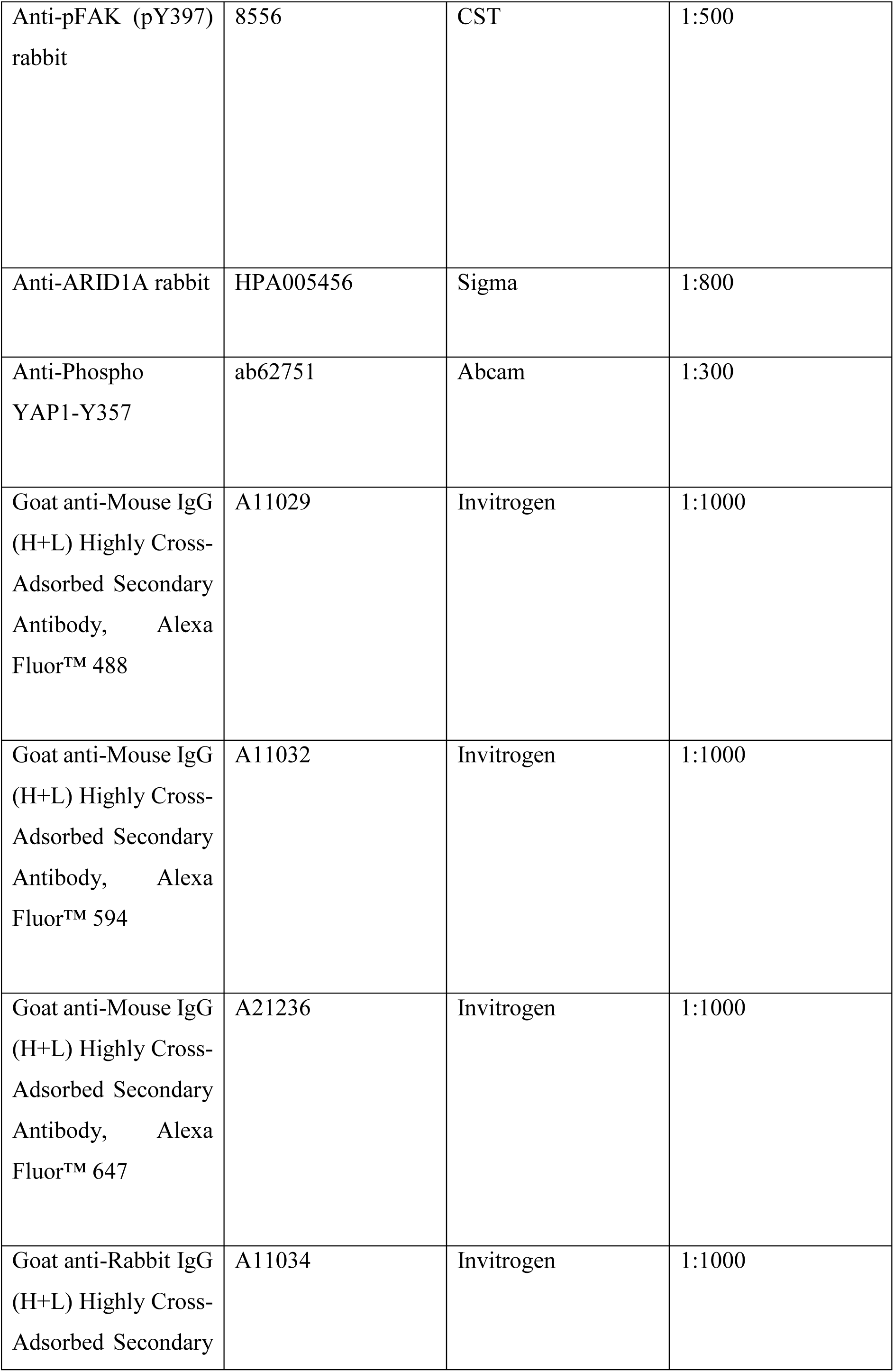
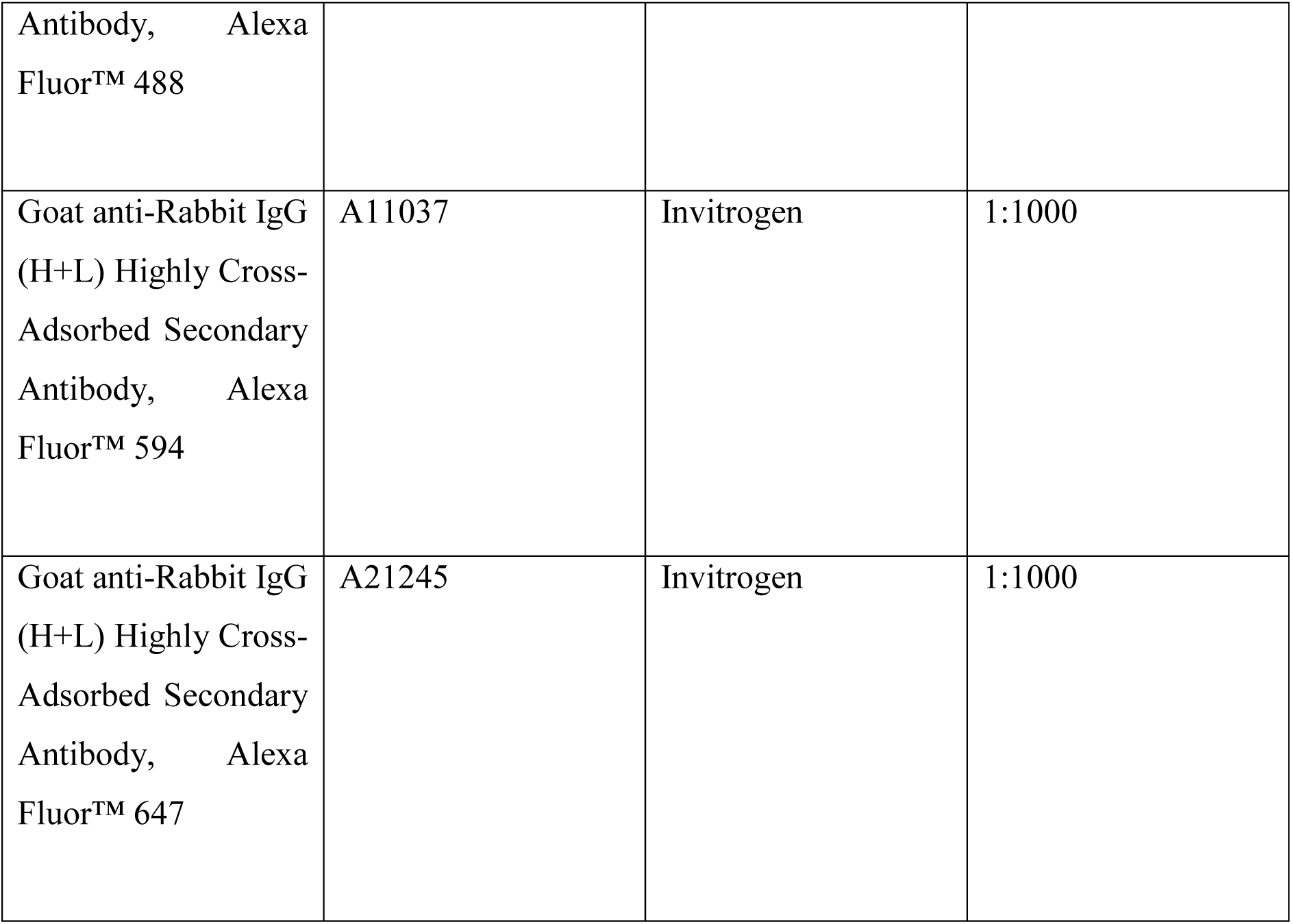

## Funding and Acknowledgements

This project was supported by intramural funds at TIFR Hyderabad from the Department of Atomic Energy, Government of India (Project Identification No. RTI 4007). We thank the Imaging and Flow Facility at TIFR, for their support access to instrumentation. We thank Prof. Tamal Das’s group for providing mApple-C1, mApple-dnNesprin1, mApple-dnLaminB1 and Jasplakinolide, and Prof. Ullas Kolthur’s group for providing the NIH/3T3 cells.

## References

1. Varelas, X., The Hippo pathway effectors TAZ and YAP in development, homeostasis and disease. Development, 2014. 141(8): p. 1614–26.

2. Zhao, B., et al., The Hippo-YAP pathway in organ size control and tumorigenesis: an updated version. Genes Dev, 2010. 24(9): p. 862–74.

3. Moroishi, T., C.G. Hansen, and K.L. Guan, The emerging roles of YAP and TAZ in cancer. Nat Rev Cancer, 2015. 15(2): p. 73–79.

4. Vassilev, A., et al., TEAD/TEF transcription factors utilize the activation domain of YAP65, a Src/Yes-associated protein localized in the cytoplasm. Genes Dev, 2001. 15(10): p. 1229–41.

5. Plouffe, S.W., A.W. Hong, and K.L. Guan, Disease implications of the Hippo/YAP pathway. Trends Mol Med, 2015. 21(4): p. 212–22.

6. Zanconato, F., M. Cordenonsi, and S. Piccolo, YAP/TAZ at the Roots of Cancer. Cancer Cell, 2016. 29(6): p. 783–803.

7. Yagi, R., et al., A WW domain-containing yes-associated protein (YAP) is a novel transcriptional co-activator. EMBO J, 1999. 18(9): p. 2551–62.

8. Strano, S., et al., Physical interaction with Yes-associated protein enhances p73 transcriptional activity. J Biol Chem, 2001. 276(18): p. 15164–73.

9. Huang, J., et al., The Hippo signaling pathway coordinately regulates cell proliferation and apoptosis by inactivating Yorkie, the Drosophila Homolog of YAP. Cell, 2005. 122(3): p. 421–34.

10. Xu, T., et al., Identifying tumor suppressors in genetic mosaics: the Drosophila lats gene encodes a putative protein kinase. Development, 1995. 121(4): p. 1053–63.

11. Harvey, K.F., C.M. Pfleger, and I.K. Hariharan, The Drosophila Mst ortholog, hippo, restricts growth and cell proliferation and promotes apoptosis. Cell, 2003. 114(4): p. 457–67.

12. Lei, Q.Y., et al., TAZ promotes cell proliferation and epithelial-mesenchymal transition and is inhibited by the hippo pathway. Mol Cell Biol, 2008. 28(7): p. 2426–36.

13. Zhao, B., et al., Inactivation of YAP oncoprotein by the Hippo pathway is involved in cell contact inhibition and tissue growth control. Genes Dev, 2007. 21(21): p. 2747–61.

14. Engler, A.J., et al., Matrix elasticity directs stem cell lineage specification. Cell, 2006. 126(4): p. 677–89.

15. Gilbert, P.M., et al., Substrate elasticity regulates skeletal muscle stem cell self-renewal in culture. Science, 2010. 329(5995): p. 1078–81.

16. Elosegui-Artola, A., et al., Mechanical regulation of a molecular clutch defines force transmission and transduction in response to matrix rigidity. Nat Cell Biol, 2016. 18(5): p. 540–8.

17. Zhang, X., et al., Talin depletion reveals independence of initial cell spreading from integrin activation and traction. Nat Cell Biol, 2008. 10(9): p. 1062–8.

18. Lombardi, M.L., et al., The interaction between nesprins and sun proteins at the nuclear envelope is critical for force transmission between the nucleus and cytoskeleton. J Biol Chem, 2011. 286(30): p. 26743–53.

19. Schwartz, M.A., Integrins and extracellular matrix in mechanotransduction. Cold Spring Harb Perspect Biol, 2010. 2(12): p. a005066.

20. Dupont, S., et al., Role of YAP/TAZ in mechanotransduction. Nature, 2011. 474(7350): p. 179–83.

21. Aragona, M., et al., A mechanical checkpoint controls multicellular growth through YAP/TAZ regulation by actin-processing factors. Cell, 2013. 154(5): p. 1047–1059.

22. Martin, K., et al., PAK proteins and YAP-1 signaling downstream of integrin beta-1 in myofibroblasts promote liver fibrosis. Nat Commun, 2016. 7: p. 12502.

23. Caliari, S.R., et al., Stiffening hydrogels for investigating the dynamics of hepatic stellate cell mechanotransduction during myofibroblast activation. Sci Rep, 2016. 6: p. 21387.

24. Elosegui-Artola, A., et al., Force Triggers YAP Nuclear Entry by Regulating Transport across Nuclear Pores. Cell, 2017. 171(6): p. 1397–1410 e14.

25. Chang, L., et al., The SWI/SNF complex is a mechanoregulated inhibitor of YAP and TAZ. Nature, 2018. 563(7730): p. 265–269.

26. Dos Santos, A., et al., DNA damage alters nuclear mechanics through chromatin reorganization. Nucleic Acids Res, 2021. 49(1): p. 340–353.

27. Shah, P., et al., ATM Modulates Nuclear Mechanics by Regulating Lamin A Levels. Front Cell Dev Biol, 2022. 10: p. 875132.

28. Kumar, A., et al., ATR mediates a checkpoint at the nuclear envelope in response to mechanical stress. Cell, 2014. 158(3): p. 633–46.

29. Basu, S., et al., Akt phosphorylates the Yes-associated protein, YAP, to induce interaction with 14-3-3 and attenuation of p73-mediated apoptosis. Mol Cell, 2003. 11(1): p. 11–23.

30. Matallanas, D., et al., RASSF1A elicits apoptosis through an MST2 pathway directing proapoptotic transcription by the p73 tumor suppressor protein. Mol Cell, 2007. 27(6): p. 962–75.

31. Hamilton, G., et al., ATM regulates a RASSF1A-dependent DNA damage response. Curr Biol, 2009. 19(23): p. 2020–5.

32. Romano, D., et al., Protein interaction switches coordinate Raf-1 and MST2/Hippo signaling. Nat Cell Biol, 2014. 16(7): p. 673–84.

33. Baskaran, R., et al., Ataxia telangiectasia mutant protein activates c-Abl tyrosine kinase in response to ionizing radiation. Nature, 1997. 387(6632): p. 516–9.

34. Shafman, T., et al., Interaction between ATM protein and c-Abl in response to DNA damage. Nature, 1997. 387(6632): p. 520–3.

35. Levy, D., et al., Yap1 phosphorylation by c-Abl is a critical step in selective activation of proapoptotic genes in response to DNA damage. Mol Cell, 2008. 29(3): p. 350–61.

36. Levy, D., et al., The Yes-associated protein 1 stabilizes p73 by preventing Itch-mediated ubiquitination of p73. Cell Death Differ, 2007. 14(4): p. 743–51.

37. Kozlov, S.V., et al., Involvement of novel autophosphorylation sites in ATM activation. EMBO J, 2006. 25(15): p. 3504–14.

38. Zhao, H. and H. Piwnica-Worms, ATR-mediated checkpoint pathways regulate phosphorylation and activation of human Chk1. Mol Cell Biol, 2001. 21(13): p. 4129–39.

39. Blackford, A.N. and S.P. Jackson, ATM, ATR, and DNA-PK: The Trinity at the Heart of the DNA Damage Response. Mol Cell, 2017. 66(6): p. 801–817.

40. Kruhlak, M.J., et al., Changes in chromatin structure and mobility in living cells at sites of DNA double-strand breaks. J Cell Biol, 2006. 172(6): p. 823–34.

41. Rogakou, E.P., et al., Megabase chromatin domains involved in DNA double-strand breaks in vivo. J Cell Biol, 1999. 146(5): p. 905–16.

42. Kobayashi, J., et al., Nucleolin participates in DNA double-strand break-induced damage response through MDC1-dependent pathway. PLoS One, 2012. 7(11): p. e49245.

43. Zhang, J., et al., YAP-dependent induction of amphiregulin identifies a non-cell-autonomous component of the Hippo pathway. Nat Cell Biol, 2009. 11(12): p. 1444–50.

44. Kudo, N., et al., Leptomycin B inhibition of signal-mediated nuclear export by direct binding to CRM1. Exp Cell Res, 1998. 242(2): p. 540–7.

45. Mazumder, A., et al., Dynamics of chromatin decondensation reveals the structural integrity of a mechanically prestressed nucleus. Biophys J, 2008. 95(6): p. 3028–35.

46. Mazumder, A. and G.V. Shivashankar, Emergence of a prestressed eukaryotic nucleus during cellular differentiation and development. J R Soc Interface, 2010. 7 Suppl 3(Suppl 3): p. S321–30.

47. Uzer, G., et al., Cell Mechanosensitivity to Extremely Low-Magnitude Signals Is Enabled by a LINCed Nucleus. Stem Cells, 2015. 33(6): p. 2063–76.

48. Bubb, M.R., et al., Effects of jasplakinolide on the kinetics of actin polymerization. An explanation for certain in vivo observations. J Biol Chem, 2000. 275(7): p. 5163–70.

49. Burma, S., et al., ATM phosphorylates histone H2AX in response to DNA double-strand breaks. J Biol Chem, 2001. 276(45): p. 42462–7.

50. Ward, I.M. and J. Chen, Histone H2AX is phosphorylated in an ATR-dependent manner in response to replicational stress. J Biol Chem, 2001. 276(51): p. 47759–62.

51. Razafsky, D. and D. Hodzic, Bringing KASH under the SUN: the many faces of nucleo-cytoskeletal connections. J Cell Biol, 2009. 186(4): p. 461–72.

52. Mellad, J.A., D.T. Warren, and C.M. Shanahan, Nesprins LINC the nucleus and cytoskeleton. Curr Opin Cell Biol, 2011. 23(1): p. 47–54.

53. Ellis, D.J., et al., GST-lamin fusion proteins act as dominant negative mutants in Xenopus egg extract and reveal the function of the lamina in DNA replication. J Cell Sci, 1997. 110 (Pt 20): p. 2507–18.

54. Solier, S. and Y. Pommier, The nuclear gamma-H2AX apoptotic ring: implications for cancers and autoimmune diseases. Cell Mol Life Sci, 2014. 71(12): p. 2289–97.

55. Toh, P.J.Y., et al., Optogenetic control of YAP cellular localization and function. EMBO Rep, 2022. 23(9): p. e54401.

56. Ota, M. and H. Sasaki, Mammalian Tead proteins regulate cell proliferation and contact inhibition as transcriptional mediators of Hippo signaling. Development, 2008. 135(24): p. 4059–69.

57. Zhao, B., et al., TEAD mediates YAP-dependent gene induction and growth control. Genes Dev, 2008. 22(14): p. 1962–71.

58. Park, J.H., et al., Mammalian SWI/SNF complexes facilitate DNA double-strand break repair by promoting gamma-H2AX induction. EMBO J, 2006. 25(17): p. 3986–97.

59. Calses, P.C., et al., TEAD Proteins Associate With DNA Repair Proteins to Facilitate Cellular Recovery From DNA Damage. Mol Cell Proteomics, 2023. 22(2): p. 100496.

60. Liaw, H., D. Lee, and K. Myung, DNA-PK-dependent RPA2 hyperphosphorylation facilitates DNA repair and suppresses sister chromatid exchange. PLoS One, 2011. 6(6): p. e21424.

61. Byrne, B.M. and G.G. Oakley, Replication protein A, the laxative that keeps DNA regular: The importance of RPA phosphorylation in maintaining genome stability. Semin Cell Dev Biol, 2019. 86: p. 112–120.

62. Olson, E., et al., RPA2 is a direct downstream target for ATR to regulate the S-phase checkpoint. J Biol Chem, 2006. 281(51): p. 39517–33.

63. Murphy, A.K., et al., Phosphorylated RPA recruits PALB2 to stalled DNA replication forks to facilitate fork recovery. J Cell Biol, 2014. 206(4): p. 493–507.

64. Pan, D., The hippo signaling pathway in development and cancer. Dev Cell, 2010. 19(4): p. 491–505.

65. Halder, G. and R.L. Johnson, Hippo signaling: growth control and beyond. Development, 2011. 138(1): p. 9–22.

66. Cordenonsi, M., et al., The Hippo transducer TAZ confers cancer stem cell-related traits on breast cancer cells. Cell, 2011. 147(4): p. 759–72.

67. Pothapragada, S.P., et al., Matrix mechanics regulates epithelial defence against cancer by tuning dynamic localization of filamin. Nat Commun, 2022. 13(1): p. 218.

68. Liang, R. and G. Song, Matrix stiffness-driven cancer progression and the targeted therapeutic strategy. Mechanobiol Med, 2023. 1(2): p. 100013.

69. Chen, B.R. and B.P. Sleckman, The regulation of DNA end resection by chromatin response to DNA double strand breaks. Front Cell Dev Biol, 2022. 10: p. 932633.

70. Bartucci, M., et al., TAZ is required for metastatic activity and chemoresistance of breast cancer stem cells. Oncogene, 2015. 34(6): p. 681–90.

71. Yu, M., et al., The mechanism of YAP/TAZ transactivation and dual targeting for cancer therapy. Nat Commun, 2025. 16(1): p. 3855.

72. Zhao, B., et al., Cell detachment activates the Hippo pathway via cytoskeleton reorganization to induce anoikis. Genes Dev, 2012. 26(1): p. 54–68.

73. Totaro, A., T. Panciera, and S. Piccolo, YAP/TAZ upstream signals and downstream responses. Nat Cell Biol, 2018. 20(8): p. 888–899.

74. Dinant, C., et al., Activation of multiple DNA repair pathways by sub-nuclear damage induction methods. J Cell Sci, 2007. 120(Pt 15): p. 2731–40.

75. Kesavan, P.S., et al., Monitoring global changes in chromatin compaction states upon localized DNA damage with tools of fluorescence anisotropy. Mol Biol Cell, 2020. 31(13): p. 1403–1410.

76. Birmingham, A., et al., 3’ UTR seed matches, but not overall identity, are associated with RNAi off-targets. Nat Methods, 2006. 3(3): p. 199–204.

77. Anderson, E.M., et al., Experimental validation of the importance of seed complement frequency to siRNA specificity. RNA, 2008. 14(5): p. 853–61.

