## Supplementary Information for "Mechanics-dependent Global Nuclear Eviction and Site-Specific Recruitment of YAP Regulates DNA Damage Responses"

### Supplementary Figures

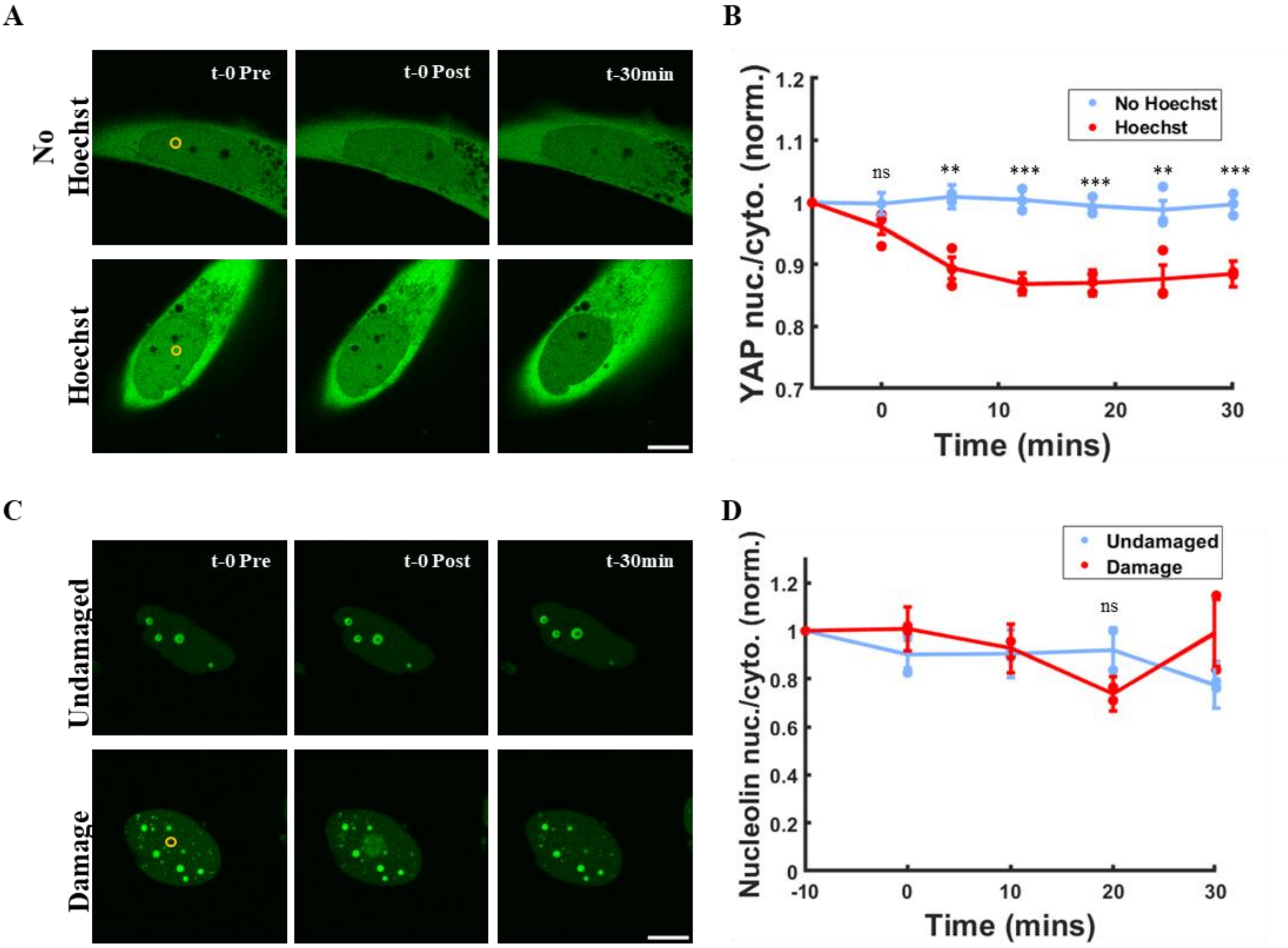


**Figure S1. YAP eviction is not due to 405nm laser caused-photobleaching and the eviction is specific to YAP**

(A) Representative images showing dynamics of EGFP-YAP1 at different time points in U2OS cells. The top panel shows non-Hoechst stained and the bottom panel shows Hoechst stained cells just before causing damage (t-0 Pre), just after damage (t-0 Post) and 30min after damage. The yellow circle represents the ROI of laser irradiation in the bottom panel. The scale bar is 10 microns.

(B) Timelapse curves showing the dynamics of EGFP-YAP1 at different time points, in non-Hoechst stained cells (blue) and Hoechst stained cells (red). The nuclear-to-cytoplasmic intensity ratio of YAP1 was measured at each time point and normalised to the initial value. Each dot represents the experiment-wise mean YAP ratio. The curve shows the overall mean ± SEM from three independent experiments (N=3, n>15 cells per condition). Statistics: The stars denote the p-values for each time point calculated using the Student's t-test ($p>0.05$-ns (non-significant), $0.01<p<0.05$ - *, $0.001<p<0.01$ - **, $p<0.001$ - ***).

(C) Representative images showing dynamics of EGFP-Nucleolin at different time points in U2OS cells. The top panel shows undamaged and the bottom panel shows cells just before causing damage (t-0 Pre), just after damage (t-0 Post) and 30min after damage. The yellow circle represents the ROI of laser irradiation in the bottom panel. The scale bar is 10 microns.

(D) Timelapse curves showing the dynamics of EGFP-Nucleolin at different time points, in undamaged cells (blue) and damaged cells (red). The nuclear-to-cytoplasmic intensity ratio of EGFP-Nucleolin was measured at each time point and normalised to the initial value. Each dot represents the experiment-wise mean EGFP-Nucleolin ratio. The curve shows the overall mean ± SEM from two independent experiments ($N=2$, $n>15$ cells per condition). Statistics: The stars denote the p-values for each time point calculated using the Student’s t-test ($p>0.05$-ns (non-significant), $0.01<p<0.05$ - *, $0.001<p<0.01$ - **, $p<0.001$ - ***).

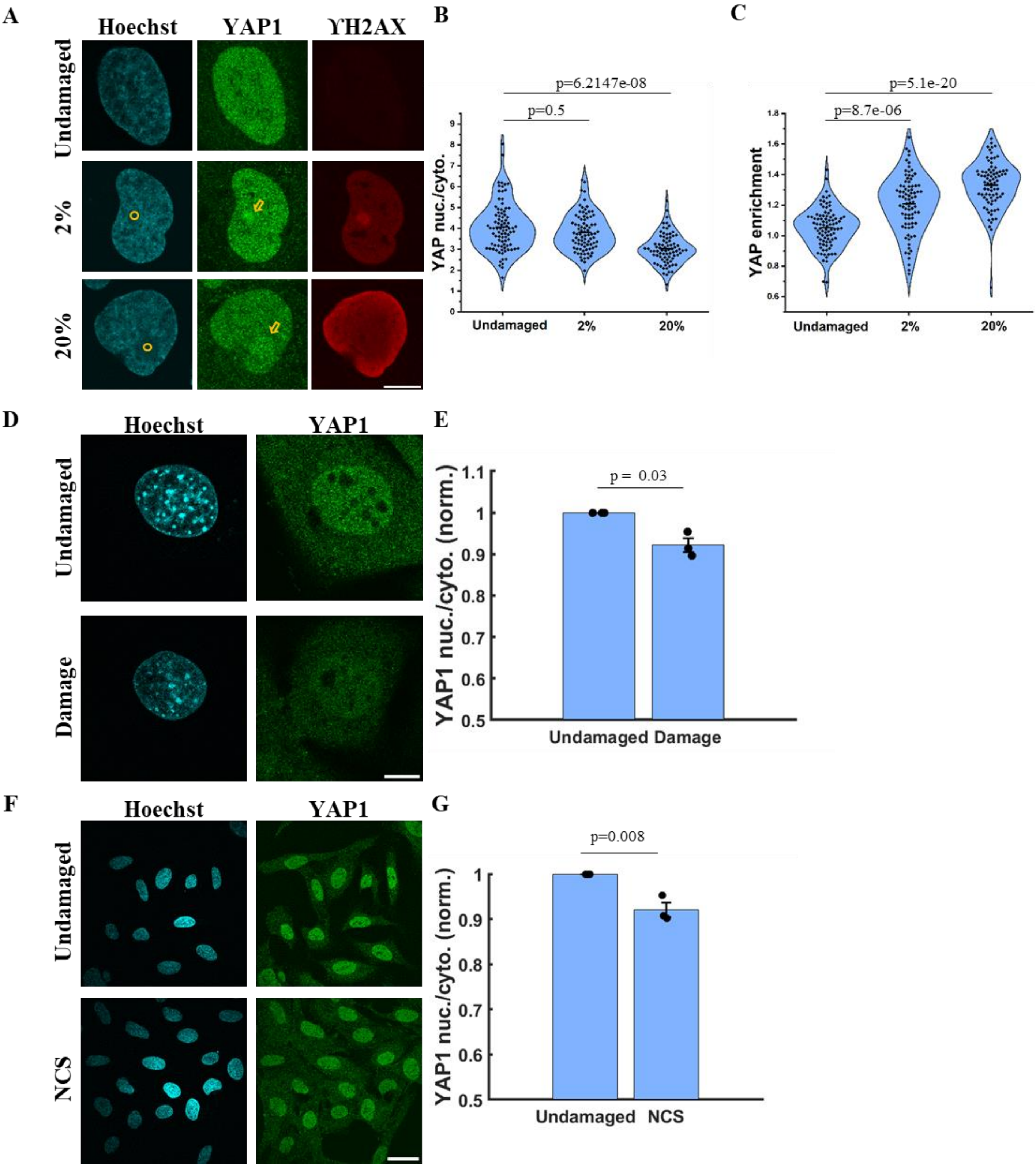


**Figure S2. DNA damage dose dependent eviction and enrichment of YAP1**

(A) Representative images showing U2OS cell nuclei with Hoechst (cyan), intensity of endogenous YAP1 levels (green) and ϒH2AX levels (red) in undamaged cells (top panel) and laser irradiated damage cells at 2% intensity (middle) and 20% intensity (bottom) of 405nm laser. The yellow circle represents the ROI of laser irradiation in the bottom panel. The yellow arrow indicates the enrichment of the proteins after damage. The scale bar is 10 microns.

(B) Violin plot with distribution quantifying nuclear-to-cytoplasmic intensity ratio of YAP1 (YAP1 nuc./cyto. or YAP ratio) for undamaged, 2% intensity and 20% intensity of 405nm laser. Each dot is the mean YAP ratio from a single cell (N=3, n>70 cells per condition). The p-values from the distributions are calculated using the Kolmogorov-Smirnov test.

(C) Violin plot with distribution quantifying YAP1 enrichment in undamaged cells, on 2% and 20% 405nm laser intensity. Each dot is the mean YAP1 enrichment from a single cell (N=3, n>70 cells per condition). Enrichment is defined as the ratio of mean intensity at the site of damage to the mean intensity in the rest of the nucleus. For undamaged cell a random spot in the nucleus for each cell was chosen from the Hoechst channel as ROI. The p-values from the distributions are calculated using the Kolmogorov-Smirnov test.

(D) Representative images showing NIH3T3 cell nuclei with Hoechst (cyan), intensity of endogenous YAP1 levels (green) in undamaged cells (top panel) and damaged cells (bottom panel). The scale bar is 10 microns.

(E) Bar graph showing YAP ratio in undamaged and damaged cells in NIH3T3 cells. Each dot represents the experiment-wise mean YAP ratio, normalised to the corresponding undamaged control, and pooled across experiments (N=3, n>70 cells per condition). The p-values are calculated using the Student’s t-test.

(F) Representative images showing U2OS cell nuclei with Hoechst (cyan), intensity of endogenous YAP1 levels (green) in undamaged cells (vehicle treated, top panel) and Neocarzinostatin (NCS) (10 ng/ml for 4 h) damaged cells (bottom panel). The scale bar is 40 microns.

(G) Bar graph showing YAP ratio in undamaged (vehicle treated) cells and NCS damaged cells. Each dot represents the experiment-wise mean YAP ratio, normalised to the corresponding undamaged control, and pooled across experiments (N=3, n>400 cells per condition). The p-values are calculated using the Student’s t-test.

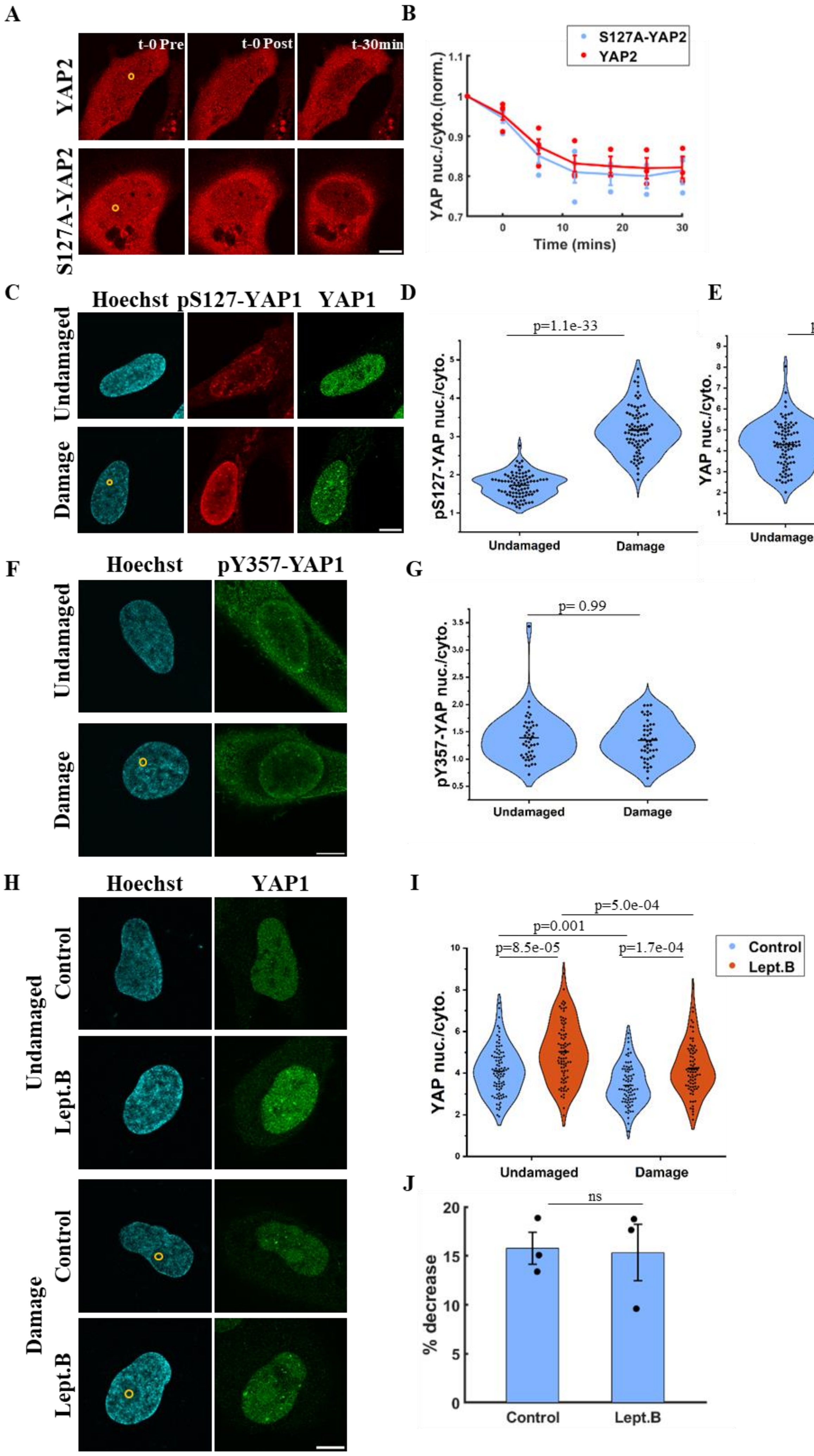
A
t-0 Pre
t-0 Post
t-30min
YAP2
S127A-YAP2
B
S127A-YAP2
YAP2
YAP nuc./cyto.(norm.)
Time (mins)
C
Hoechst
pS127-YAP1
YAP1
Undamaged
Damage
D
p=1.1e-33
pS127-YAP nuc./cyto.
Undamaged
Damage
E
p=2.8e-05
YAP nuc./cyto.
Undamaged
Damage
F
Hoechst
pY357-YAP1
Undamaged
Damage
G
p= 0.99
pY357-YAP nuc./cyto.
Undamaged
Damage
H
Hoechst
YAP1
Undamaged
Control
Lept.B
Damage
Control
Lept.B
I
p=5.0e-04
p=0.001
p=8.5e-05
p=1.7e-04
Control
Lept.B
YAP nuc./cyto.
Undamaged
Damage
J
ns
% decrease
Control
Lept.B

**Figure S3. DNA damage driven YAP nuclear eviction is independent of Hippo pathway**

(A) Representative images showing dynamics of dsRed-YAP2 and dsRed-S127A-YAP2 overexpressing U2OS cells at different time points. The top panel shows dsRed-YAP2 cells and the bottom panel shows dsRed-S127A-YAP2 cells just before causing damage (t-0 Pre), just after damage (t-0 Post) and 30min after damage. The yellow circle represents the ROI of laser irradiation. The scale bar is 10 microns.

(B) Timelapse curves showing the dynamics of dsRed-YAP2 and dsRed-S127A-YAP2 at different time points, in undamaged control (blue) and damage cells (red). The nuclear-to-cytoplasmic intensity ratio of YAP1 was measured at each time point and normalised to the initial value. Each dot represents the mean normalised value from one experiment. The curve shows the overall mean ± SEM from three independent experiments (N=3, >20 cells per condition). Statistics: The stars denote the p-values for each time point calculated using the Student's t-test (p>0.05-ns, 0.01<p<0.05 - *, 0.001<p<0.01 - **, p<0.001 - ***).

(C) Representative images showing U2OS cell nuclei with Hoechst (cyan), intensity of endogenous pS127-YAP1 (red) and YAP1 (green) in undamaged cells (top panel) and damage cells (bottom panel). The yellow circle represents the ROI of laser irradiation in the bottom panel. The scale bar is 10 microns.

(D) Violin plot with distribution quantifying nuclear-to-cytoplasmic intensity ratio of pS127-YAP1 (pS127-YAP1 nuc./cyto.) in undamaged and damaged cells. Each dot is the mean pS127-YAP1 nuc./cyto. from a single cell (N=3, n>70 cells per condition). The p-values from the distributions is calculated using the Kolmogorov-Smirnov test.

(E) Violin plot with distribution quantifying nuclear-to-cytoplasmic intensity ratio of YAP1 (YAP1 nuc./cyto. or YAP ratio) in undamaged and damage cells. Each dot is the mean YAP1 nuc./cyto. from a single cell (N=3, n>70 cells per condition). The p values from the distributions is calculated using the Kolmogorov-Smirnov test.

(F) Representative images showing U2OS cell nuclei with Hoechst (cyan), intensity of endogenous pY357-YAP1 (green) in undamaged cells (top panel) and damage cells (bottom panel). The yellow circle represents the ROI of laser irradiation in the bottom panel. The scale bar is 10 microns.

(G) Violin plot with distribution quantifying nuclear-to-cytoplasmic intensity ratio of pY357-YAP1 (pY357-YAP1 nuc./cyto.) in undamaged and damaged cells. Each dot is the mean pY357-YAP1 nuc./cyto. from a single cell (N=2, n>50 cells per condition). The p-values from the distributions is calculated using the Kolmogorov-Smirnov test.

(H) Representative images showing U2OS cell nuclei with Hoechst (cyan), intensity of endogenous YAP1 levels (green) in undamaged cells (top panel) and damage cells (bottom panel) in both vehicle treated (Control, DMSO) and Leptomycin B treated (Lept.B) cells. The yellow circle represents the ROI of laser irradiation in the bottom panel. The scale bar is 10 microns.

(I) Violin plot with distribution quantifying nuclear-to-cytoplasmic intensity ratio of YAP1 (YAP1 nuc./cyto. or YAP ratio) in undamaged and damaged cells of both vehicle treated (Control, blue) and Leptomycin B treated (Lept.B, orange) cells. Each dot is the mean YAP ratio from a single cell (N=3, n>70 cells per condition). The p-values from the distributions is calculated using the Kolmogorov-Smirnov test.

(J) Bar graph showing the percentage decrease in YAP ratio in Control cells and Lept.B treated cells on DNA damage. Each dot represents the mean % decrease obtained from each experiment ($N=3$, $n>70$ cells per condition). The p-values from the distributions is calculated using the Student's t-test ($p>0.05$-ns, $0.01<p<0.05$ - *, $0.001<p<0.01$ - **, $p<0.001$ - ***).

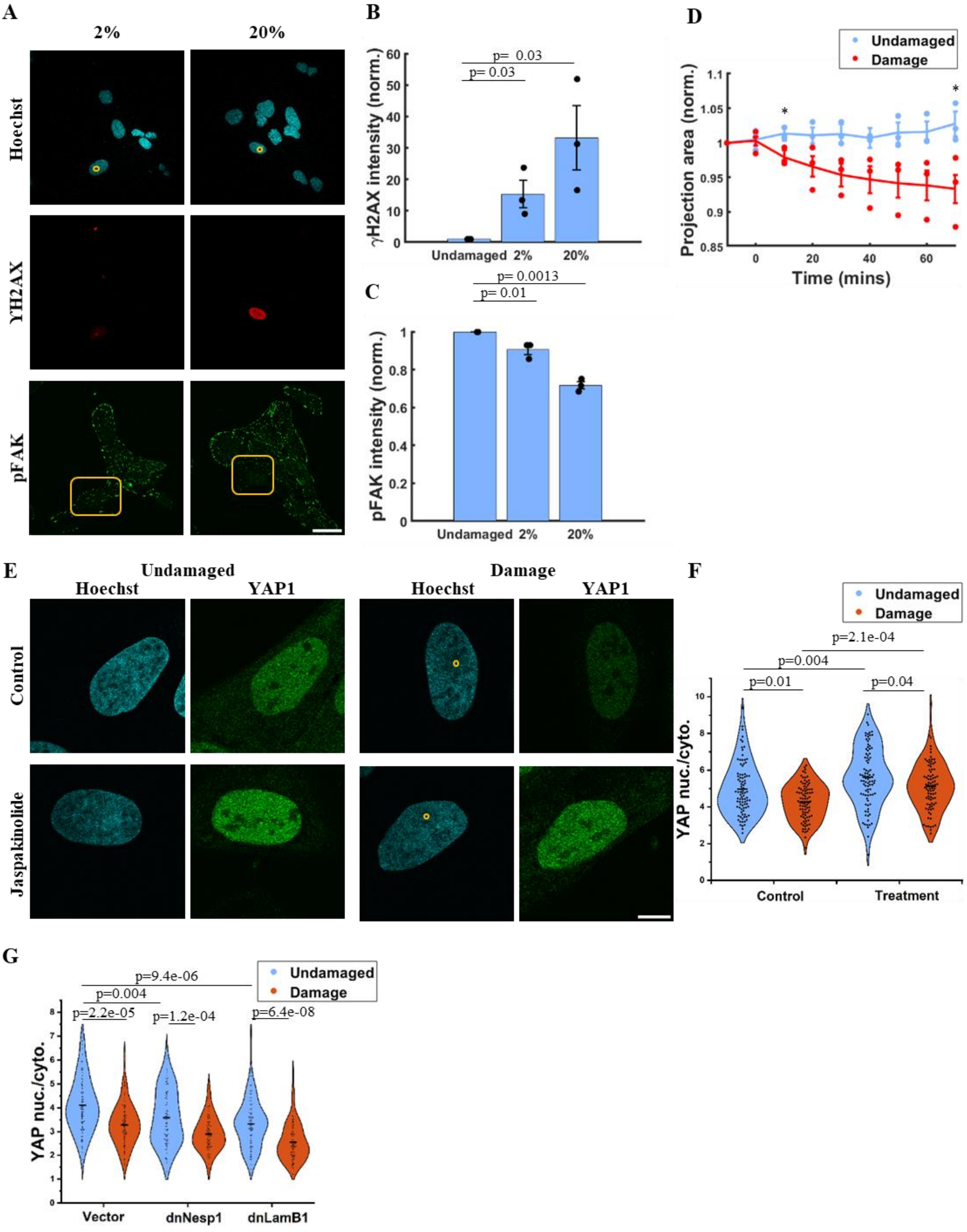

A
2%
20%
Hoechst
γH2AX
pFAK
B
γH2AX intensity (norm.)
p= 0.03
p= 0.03
Undamaged 2% 20%
C
p= 0.0013
p= 0.01
pFAK intensity (norm.)
Undamaged 2% 20%
D
Undamaged
Damage
Projection area (norm.)
Time (mins)
E
Undamaged
Damage
Hoechst
YAP1
Control
Jaspakinolide
F
Undamaged
Damage
p=2.1e-04
p=0.004
p=0.01
p=0.04
YAP nuc./cyto.
Control
Treatment
G
Undamaged
Damage
p=9.4e-06
p=0.004
p=2.2e-05
p=1.2e-04
p=6.4e-08
YAP nuc./cyto.
Vector
dnNesp1
dnLamB1

**Figure S4. Nuclear mechanics regulates YAP nuclear levels**

(A) Representative images showing U2OS cell nuclei with Hoechst (cyan), endogenous ϒH2AX (red) and pFak (pY397) (green) on 2% and 20% 405nm laser intensity. The yellow circle represents the ROI of laser irradiation in the top panel. The scale bar is 40 microns.

(B) Bar graphs quantifying ϒH2AX intensity in undamaged cells, on 2% and 20% 405nm laser damage. Each dot represents the experiment-wise mean, normalised to the corresponding undamaged control, and pooled across experiments (N=3, n>70 cells per condition). The p-values from the distributions are calculated using unpaired Student's t-test.

(C) Bar graphs with distribution quantifying total intensity of pFAK at focal adhesions in undamaged, 2% and 20% 405nm laser damaged cells. Each dot represents the experiment-wise mean, normalised to the corresponding undamaged control, and pooled across experiments (N=3, n>70 cells per condition). The p-values from the distributions are calculated using unpaired Student's t-test.

(D) Timelapse curves showing the change in nuclear projection area of EGFP-YAP1 overexpressing cells at different time points in undamaged (blue) and damaged cells (red). The projection area was measured at each time point and normalised to the initial projection area before damage induction. Each dot represents the experiment-wise mean normalised value. The curve shows the overall mean ± SEM from three independent experiments (N=3, >20 cells per condition). Statistics: The stars denote the p-values for each time point calculated using the student's t-test (p>0.05-ns, 0.01<p<0.05 - *, 0.001<p<0.01 - **, p<0.001 - ***).

(E) Representative images showing U2OS cell nuclei with Hoechst (cyan), intensity of endogenous YAP1 levels (green) in undamaged cells (left panel) and damage cells (right panel) in both vehicle treated (Control, DMSO) and Jaspakinolide treated cells. The yellow circle represents the ROI of laser irradiation in the right panel. The scale bar is 10 microns.

(F) Violin plot with distribution quantifying nuclear-to-cytoplasmic intensity ratio of YAP1 (YAP1 nuc./cyto. or YAP ratio) in undamaged (blue) and damaged (orange) cells of both vehicle treated (Control) and Jasplakinolide treated (Treatment) cells. Each dot is the mean YAP ratio from a single cell (N=3, n>70 cells per condition). The p-values from the distributions is calculated using the Kolmogorov-Smirnov test.

(G) Violin plot with distribution quantifying YAP ratio in undamaged (blue) and damaged (orange) cells of vector, dnNesp1 and dnLamB1 overexpressing U2OS cells (N=3, n>70 cells per condition). Each dot is the YAP ratio from a single cell. The p-values from the distributions are calculated using the Kolmogorov-Smirnov test.

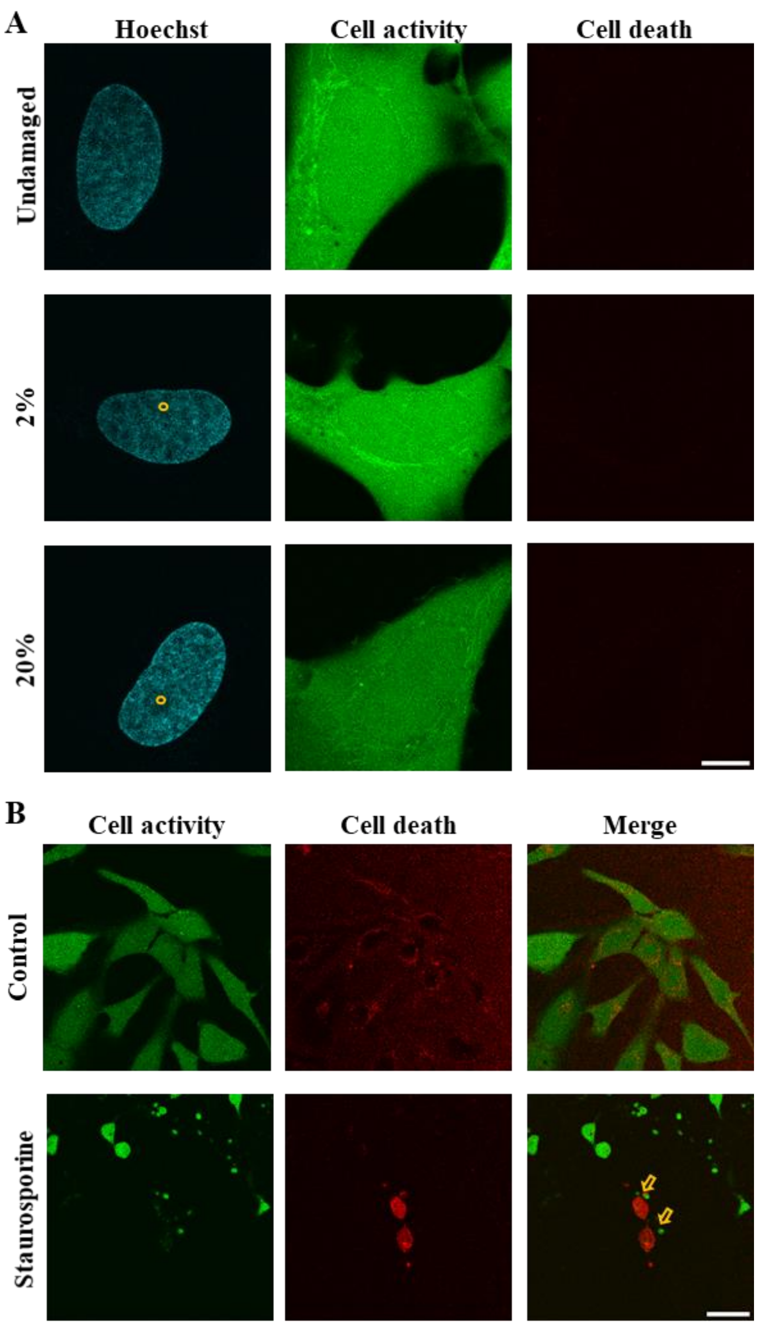


**Figure S5 Cell death is not induced on laser-induced DNA damage**

A) Representative images showing U2OS cell nuclei with Hoechst (cyan), cell activity indicator (green) or cell death indicator (red) after staining with LIVE/DEAD cell imaging kit in undamaged (top), 2% (middle) and 20% (bottom) 405nm laser irradiated cells. The yellow circle represents the ROI of laser irradiation. The scale bar is 10 microns.

B) Representative images showing U2OS cells with cell activity indicator (green) or cell death indicator (red) imaged at 60X magnification after staining with LIVE/DEAD cell imaging kit in vehicle treated (Control) and Staurosporine treated cells. The yellow arrow indicates the cells that show the intensity of the cell death stain. The scale bar is 40 microns.

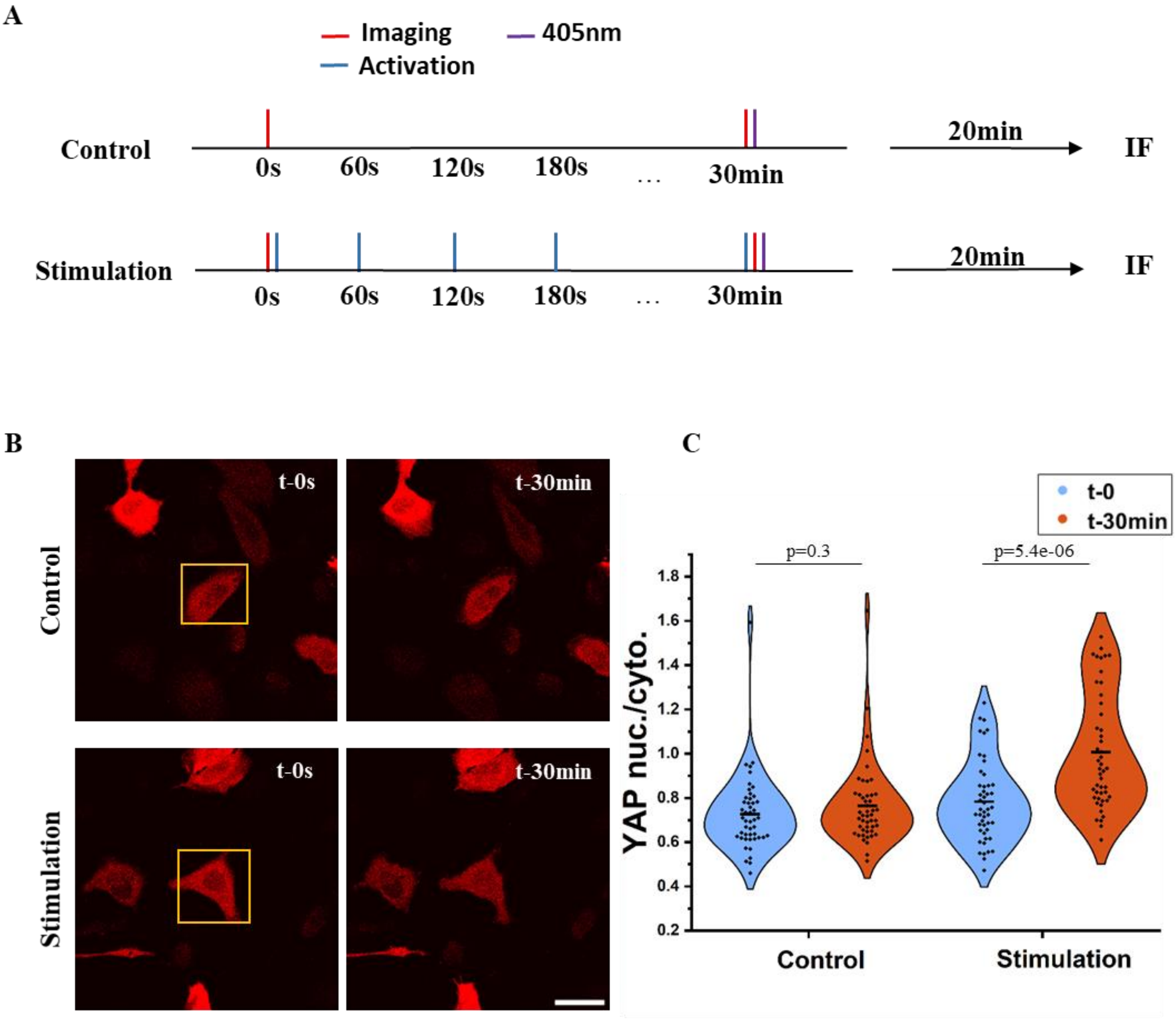


**Figure S6. Changes in YAP localisation on light activation of OptoYAP**

A) Schematic for OptoYAP stimulation followed by DNA damage. Cells initially showing higher cytoplasmic YAP localization were stimulated with 488nm laser every 60s for 30min (activation) and then laser irradiated at a point with 405nm laser (Stimulation). Unstimulated cells were also laser irradiated with 405nm (Control). 20mins after 405nm irradiation, all cells were fixed and stained for the required proteins.

B) Representative images showing intensity of OptoYAP at t-0 s and t-30 mins post activation with 488nm laser, in live control or stimulated U2OS cells. The yellow box indicates the region of interest where 488nm laser stimulation was caused. The scale bar is 10 microns.

C) Violin plot with distribution quantifying nuclear to cytoplasmic levels of YAP (YAP nuc./cyto. or YAP ratio) in live control and stimulated cells at t-0 (blue) and t-30min (orange). Each dot is the YAP ratio from a single cell. The p-values from the distributions is calculated using Kolmogorov-Smirnov test.

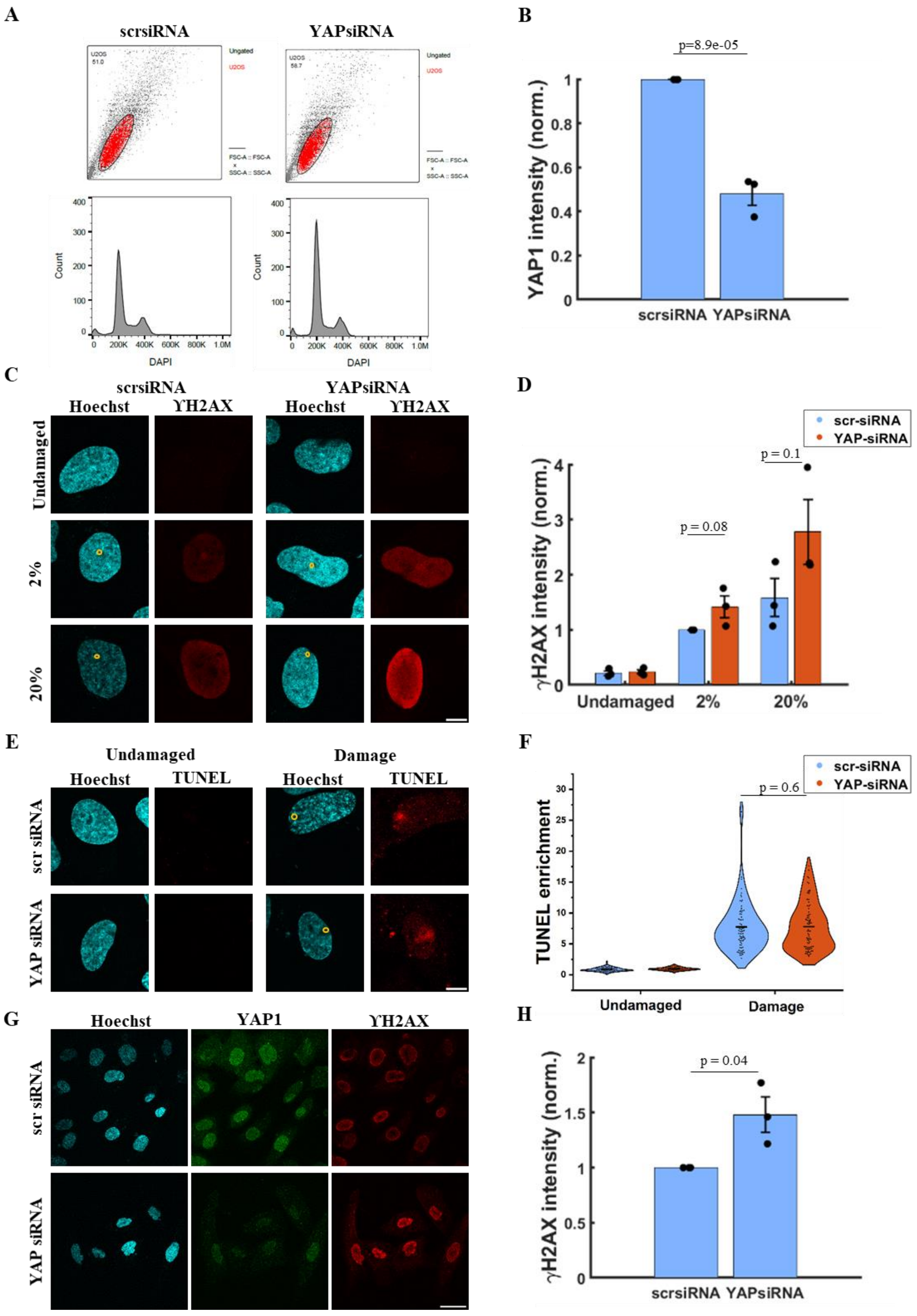
A
scrsiRNA
YAPsiRNA
Ungated
U2OS
FSC-A :: FSC-A
x
SSC-A :: SSC-A
Count
DAPI
B
p=8.9e-05
YAP1 intensity (norm.)
scrsiRNA YAPsiRNA
C
scrsiRNA
YAPsiRNA
Hoechst
ϒH2AX
Undamaged
2%
20%
D
scr-siRNA
YAP-siRNA
p = 0.1
p = 0.08
γH2AX intensity (norm.)
Undamaged
2%
20%
E
Undamaged
Damage
Hoechst
TUNEL
scr siRNA
YAP siRNA
F
scr-siRNA
YAP-siRNA
p = 0.6
TUNEL enrichment
Undamaged
Damage
G
Hoechst
YAP1
ϒH2AX
scr siRNA
YAP siRNA
H
p = 0.04
γH2AX intensity (norm.)
scrsiRNA YAPsiRNA

**Figure S7. Effect of knockdown of YAP on extent of DNA damage and DNA damage responses**

(A) DNA content analysis of U2OS cells (scrsiRNA treated and YAP siRNA treated) analyzed with the flow-cytometer (Cytoflex-S).

(B) Bar graphs quantifying nuclear intensity of YAP1 (YAP1 intensity) in undamaged scrambled (scr) or YAP siRNA treated cells. Each dot represents the experiment-wise mean, normalised to the corresponding undamaged scr-siRNA control, and pooled across experiments (N=3, n>70 cells per condition). The p-values from the distributions are calculated using unpaired Student's t-test.

(C) Representative images showing U2OS cell nuclei with Hoechst (cyan), intensity of endogenous ϒH2AX (red) in undamaged cells (top panel), 2% (middle panel) and 20% 405nm laser intensity (bottom panel) in scr-siRNA and YAP-siRNA treated cells. The yellow circle represents the ROI of laser irradiation. Scale bar- 10µm.

(D) Bar graph with distribution quantifying ϒH2AX intensity in undamaged cells, on 2% and 20% 405nm laser damaged in scr-siRNA (blue) and YAP-siRNA (orange) treated cells. Each dot represents the experiment-wise mean, normalised to the 2% scr-siRNA damage condition, and pooled across experiments (N=3, n>70 cells per condition). The p-values from the distributions are calculated using unpaired Student's t-test.

(E) Representative images showing U2OS cell nuclei with Hoechst (cyan), intensity of TUNEL assay (red) in undamaged cells (left panel) and damaged cells (right panel) in scr-siRNA and YAP-siRNA treated cells. The yellow circle represents the ROI of laser irradiation. The scale bar is 10 microns.

(F) Violin plot with distribution quantifying TUNEL enrichment in undamaged cells and damage cells on 405nm laser damage in scr-siRNA (blue) and YAP-siRNA (orange) treated cells. Each dot is the enrichment from a single cell. Enrichment is defined as the ratio of mean intensity at the site of damage to the mean intensity in the rest of the nucleus. (N=3, n>80 cells per condition). For undamaged cell a random spot in the nucleus for each cell was chosen from the Hoechst channel as ROI. The p-values from the distributions is calculated using Kolmogorov-Smirnov test.

(G) Representative images showing U2OS cell nuclei with Hoechst (cyan), intensity of endogenous YAP1(green) and ϒH2AX intensity (red) in cells of scr-siRNA (top panel) and YAP-siRNA treated cells (bottom panel). DNA damage was caused using Neocarzinostatin (NCS) for 1hr at 1µg/ml concentration. Scale bar- 40µm.

(H) Bar graph quantifying ϒH2AX intensity in scr-siRNA and YAP-siRNA treated cells damaged with NCS. Each dot represents the experiment-wise mean, normalised to the scr-siRNA control, and pooled across experiments (N=3, n>400 cells per condition). The p-values from the distributions are calculated using unpaired Student's t-test.

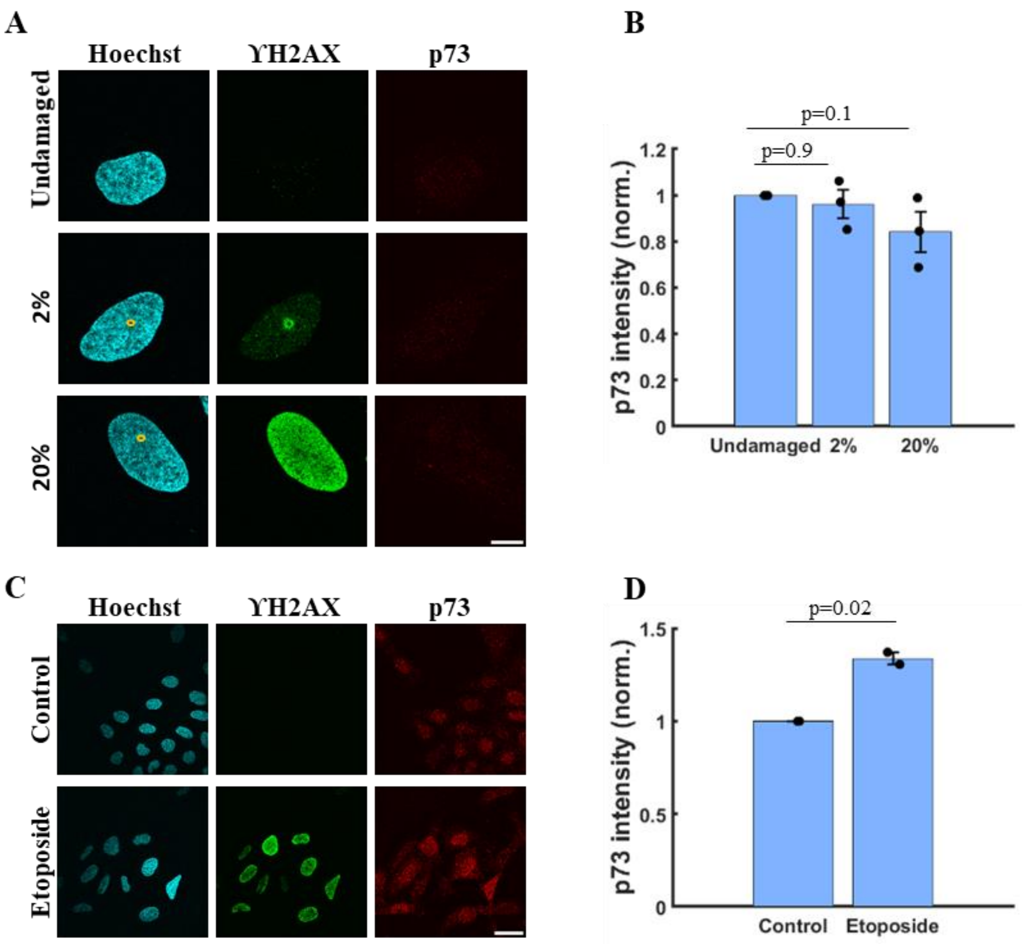


**Figure S8. p73 is not induced upon laser-induced DNA damage**

A) Representative images showing U2OS cell nuclei with Hoechst (cyan), intensity of endogenous ϒH2AX (green) and p73 (red) in undamaged cells (top panel), on 2% (middle panel) and 20% 405nm laser intensity (bottom panel). The yellow circle represents the ROI of laser irradiation. The scale bar is 10 microns.

B) Bar graphs quantifying intensity of p73 in undamaged, on 2% and 20% 405nm laser intensity. Each dot represents the experiment-wise mean, normalised to the corresponding undamaged control, and pooled across experiments (N=3, n>70 cells per condition). The p-values from the distributions are calculated using unpaired Student's t-test.

C) Representative images showing U2OS cell nuclei with Hoechst (cyan), intensity of endogenous ϒH2AX (green) and p73 (red) in vehicle treated cells (Control, DMSO, top panel) and on Etoposide treatment (bottom panel). The images were captured at 60x magnification. The scale bar is 40 microns.

D) Bar graph quantifying intensity of p73 in vehicle treated (Control, DMSO) and on Etoposide treatment. Each dot represents the experiment-wise mean, normalised to the corresponding undamaged control, and pooled across experiments (N=2, n>200 cells per condition). The p-values from the distributions are calculated using unpaired Student's t-test.

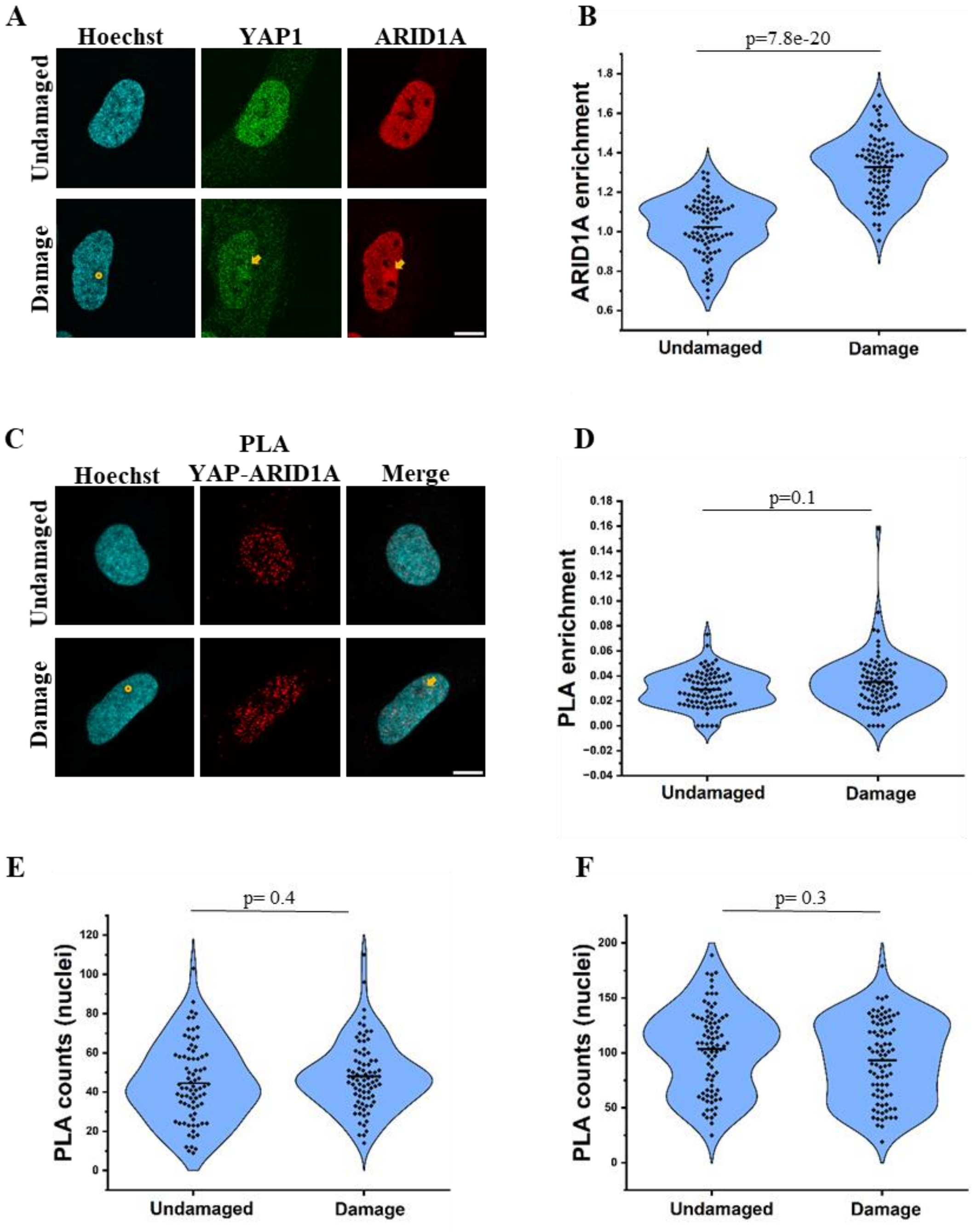


**Figure S9. ARID1A is enriched but does not interact with YAP at DNA damage site**

A) Representative images showing U2OS cell nuclei with Hoechst (cyan), intensity of endogenous YAP1 (green) and ARID1A intensity (red) in undamaged (top panel) and damaged (bottom panel) cells. The yellow circle represents the ROI of laser irradiation. The yellow arrow indicates the enrichment of the proteins after damage. The scale bar is 10 microns.

B) Violin plot with distribution quantifying enrichment of ARID1A intensity in undamaged and damaged cells. Each dot is the enrichment from a single cell (N=3, n>80 cells per condition). Enrichment is defined as the ratio of mean intensity at the site of damage to the mean intensity in the rest of the nucleus. The p-values from the distributions is calculated using the Kolmogorov-Smirnov test.

C) Representative images showing U2OS cell nuclei with Hoechst (cyan), PLA intensity of YAP1-ARID1A (red), and merge of the two channels in undamaged cells (top panel) and damaged cells (bottom panel). The yellow circle represents the ROI of laser irradiation. The yellow arrow indicates the site of PLA signal at DNA damage site. The scale bar is 10 microns.

D) Violin plot with distribution quantifying enrichment of YAP1-ARD1A PLA counts in undamaged and damaged cells. Each dot is the enrichment from a single cell (N=3, n>70 cells per condition). Enrichment is defined as the total number of PLA foci at the site of damage to the total number of PLA foci in the rest of the nucleus. For undamaged cell a random spot in the nucleus for each cell was chosen from the Hoechst channel as control. The p-values from the distributions are calculated using the Kolmogorov-Smirnov test.

E) Violin plot with distribution quantifying total nuclear PLA counts for YAP1-TEAD1 in undamaged and damaged cells. Each dot is the total nuclear PLA counts from a single cell (N=3, n>70 cells per condition). The p values from the distributions is calculated using the Kolmogorov-Smirnov test.

F) Violin plot with distribution quantifying total nuclear PLA counts for YAP1-ARID1A in undamaged and damaged cells. Each dot is the total nuclear PLA counts from a single cell (N=3, n>70 cells per condition). The p values from the distributions is calculated using the Kolmogorov-Smirnov test.

**A**

| Protein name | Gene name |
|---|---|
| Serine/threonine-protein kinase Chk1 | CHEK1 |
| Replication protein A 32KDa subunit | RPA2 |
| Double-strand-break repain protein rad21 homolog | RAD21 |
| DNA excision repair protein ERCC-6-like | ERCC6L |
| DNA-directed RNA polymerases 1,2 and 3 subunit RPABC3 | POLR2H |
| KIF1-binding protein | KIAA1279 |
| Replication factor C subunit 1 | RFC1;LLDBP |
| Actin-related protein 2/3 complex subunit 1A | ARPC1A |

Adapted from Chang et.al.

**B**

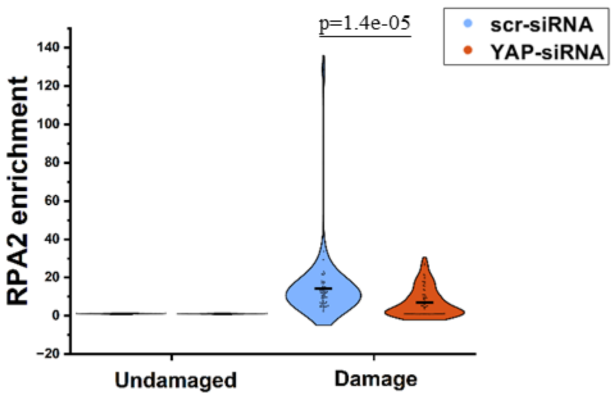


**Figure S10. Candidate YAP interactors in DDR and reduced long-term RPA2 site recruitment upon YAP knockdown**

(A) Network of putative YAP interactors associated with DNA damage response pathways. Proteins were shortlisted from a published YAP study looking at YAP interactors.

(B) Violin plot with distribution quantifying RPA2 enrichment in undamaged and damaged cells in both scrambled siRNA treated (scr-siRNA, orange) and YAP siRNA treated (YAP-siRNA, orange) cells, 6 hrs after DNA damage. Each dot is the enrichment from a single cell (N=2, n>40 cells per condition). Enrichment is defined as the ratio of mean intensity at the site of damage to the mean intensity in the rest of the nucleus. For undamaged cell a random spot in the nucleus for each cell was chosen from the Hoechst channel as ROI. The p-values from the distributions are calculated using the Kolmogorov-Smirnov test.
